# The murine intestinal pathobiont *Helicobacter hepaticus* attenuates DSS colitis in a CD4+ T cell-dependent manner

**DOI:** 10.1101/2025.10.29.685394

**Authors:** Anna L. L. Heawood, Anna A. M. Ahlback, Annika Frede, Xiang Li, David McGuinness, John Cole, Tshandyemwene R. Tshivolo, Holly C. Webster, Kevin J. Maloy

## Abstract

The host immune system is fundamentally shaped by its resident microbiota; however, much remains unknown about how individual microbial species contribute to health and disease. *Helicobacter hepaticus* (*Hh*) is a member of the murine intestinal microbiota associated with colitis in immunodeficient mice, despite driving dominant immune regulatory responses in normal hosts which allow colonization without pathology. However, whether this colonization influences intestinal immune homeostasis more widely remains unexplored. Here, we report that *Hh* colonization confers a disease protective effect on DSS colitis, attenuating intestinal inflammation and other disease parameters. Disease attenuation required persistent colonization and was dependent on host CD4^+^ T cells. We further show that *Hh* colonization promotes a conserved anti-inflammatory transcriptional programme across several effector and regulatory intestinal CD4^+^ T cell subsets. Thus, although persistent stimulation of host immune responses allows the host to tolerate ‘pathobionts’ like *Hh*, this is compensated by the promotion of tissue-protective immunological conditioning.

## Introduction

The mammalian immune system evolved alongside a complex microbiota, driving constant immune responses at steady state which occur in the absence of inflammation and help to regulate immunological function [1]. While perturbations to the intestinal microbiota have been implicated in the development of various diseases, most notably inflammatory bowel disease (IBD) [2, 3], the health-promoting effects of commensal microbes are a focus of public interest. Indeed, much research has attempted to identify specific bacterial species and families which drive either a disease-protective or worsening effect in different settings [4, 5]. However, the nature of such interactions and the mechanisms behind them are not fully understood.

Pathobionts are microbes which generally behave as commensals but have the capacity to drive inflammation in certain contexts. These species actively engage the immune system and alter intestinal homeostasis in a context-dependent manner, tending to drive pathology in the absence of key immune regulatory elements [6]. One well-characterised example is *Helicobacter hepaticus* (*Hh*), a gram-negative bacterium which colonizes the murine large intestine [7, 8]. This microbe has been described to drive colitis in certain lymphopenic mouse strains, including severe combined immunodeficiency (SCID) and recombination activating gene (RAG) ^-/-^ mice, [9-11]. Despite this, immunocompetent animals show no pathology when colonized with *Hh*, as immune regulation restrains pathogenic inflammatory responses towards the bacteria [12]. As such, the interleukin-10 (IL-10) pathway plays a critical role in the ability of mice to tolerate *Hh* infection as defects in this pathway cause colitis in *Hh* colonized mice [12-14]. Characterisation of the colonic T cell compartment has revealed that colonization preferentially drives the differentiation of *Hh*-specific regulatory T cells (Tregs) at steady state, while loss of IL-10 signalling instead results in expansion of Th1 and Th17 effectors [13-15]. Similarly, in susceptible mice, macrophages are a key cell type contributing to pathology during *Hh*-induced colitis [16, 17] – yet these cells upregulate an anti-inflammatory transcriptional program following *Hh* infection of immunocompetent animals [18]. These findings highlight the dominant regulatory effect of *Hh* on local immune phenotypes, although the outcome of these regulatory responses for subsequent host immune function remains largely unexplored.

Dextran sulphate sodium (DSS) induced colitis is a widely used rodent model of IBD, in which the administration of DSS in drinking water causes colonic epithelial injury. Loss of colonic barrier integrity results in pathological inflammatory responses targeted against the microbiota, driving an acute colitis [19]. Notably, DSS induces colitis in SCID mice lacking T and B cells, demonstrating the ability of the innate immune system to drive pathology in this model [20]. The microbiota has also been shown to play a key role in regulating the severity of DSS colitis, with specific bacterial species associated with either disease protection or aggravation [5]. Consequently, cohorts of mice with identical genetic backgrounds show large scale variation in their responsiveness to DSS [5], while differences in microbiota composition between transgenic and wild-type colonies can confound evaluations into the effect of genotype on disease severity [21]. Despite this, the mechanistic basis of how individual microbes modulate disease severity in this model remains poorly understood.

While it is clear that pathobiont species such as *Hh* exert a range of effects on host immunity, recent work comparing the microbiome of wild mice to that of laboratory mice demonstrated that these species are generally underrepresented in laboratory animals compared to wild counterparts [22]. For example, wild mice have far higher relative abundance of Proteobacteria and Helicobacteraceae compared to conventional laboratory animals, as well as a general increase in microbial diversity [22, 23]. This vast difference in microbial content is thought to drive differences in immune function, with mice bearing a wild microbiota showing a survival advantage over laboratory mice in models of lethal influenza infection and colitis-associated tumorigenesis [23]. Importantly, wild mice are thought to more accurately model human immune responses, with laboratory counterparts described to possess an immature immune system due to their reduced microbial load [22, 24, 25]. Given the high prevalence of *Helicobacter* species within the wild mouse microbiota [22] and the survival advantage this conferred in certain contexts, as well as the documented immunomodulatory effects of *Hh*, we hypothesised that *Hh* could be a candidate pathobiont species contributing to the disease attenuating effects of a wild microbiota.

Here, we explored the effects of *Hh* colonization on inflammatory disease in the context of DSS colitis. We demonstrate that infection with this microbe attenuates DSS disease severity, including alleviating clinical colitis symptoms such as weight loss and attenuating histopathological inflammation. We find that during DSS colitis, *Hh* colonized mice show markedly reduced colonic inflammatory infiltrate, despite showing increased myeloid recruitment to the colon at steady state. We further show that local macrophage phenotypes are altered following *Hh* colonization, generally exhibiting reduced inflammatory cytokine production. Finally, we show that *Hh*- mediated attenuation of DSS disease severity is dependent on CD4^+^ T cells and that *Hh* colonization enhances regulatory phenotypes across many CD4^+^ T cell subsets. Thus, while *Hh* has largely been studied as a model for pathobiont-induced colitis in susceptible mice, we demonstrate that, in immunocompetent animals, *Hh* colonization provides a beneficial function for the host by protecting against acute inflammatory challenge in the gut.

## Results

### *Hh* attenuates DSS-driven intestinal inflammation

To test whether *Hh* colonization alters the severity of intestinal inflammatory disease in immunocompetent animals, we utilized the DSS colitis model which is known to be influenced by microbiota composition [5]. Specific pathogen free (SPF) C57BL/6 mice were infected with *Hh* strain by oral gavage and left for 21 days to allow the bacteria to fully colonize (Figure 1A). Mice were then treated with 2% DSS in drinking water for 4 days before being returned to normal drinking water until the experiment endpoint 4 days later (day 29 after *Hh* colonization), while control animals remained on normal drinking water throughout (Figure 1A). In this model, DSS causes acute colonic inflammation driven primarily by myeloid cells, particularly infiltrating monocytes and neutrophils [26]. DSS-treated mice were assessed daily for clinical symptoms, including stool consistency, rectal bleeding, general appearance and weight loss, and we found that *Hh* colonized mice had significantly reduced disease activity (Figure 1B). DSS treatment caused substantial weight loss in uninfected animals, with a mean weight loss of ∼10% of starting weight (Figure 1C), whereas mice infected with *Hh* were protected against severe weight loss, with many returning almost to their starting weight by the end of the experiment (Figure 1C). We measured colon length as an indicator of intestinal inflammation, as inflammation causes a thickening of the colon tissue which results in shortening of total length [27]. We observed reduced colon lengths in both DSS-treated groups, although to a significantly lesser extent in *Hh* colonized animals (Figure 1D). Finally, we carried out histological assessment of the mid and distal colon for intestinal pathology and found that both DSS-treated groups contained samples showing cardinal signs of inflammation, characterised by epithelial hyperplasia, leukocyte infiltration, loss of crypt structure, and oedema (Figure 1E) [28, 29]. However, the *Hh* colonized group had a significantly decreased histology score compared to the PBS-treated group, indicating that *Hh* colonization attenuated DSS-induced intestinal pathology (Figure 1E).

**Figure 1:**
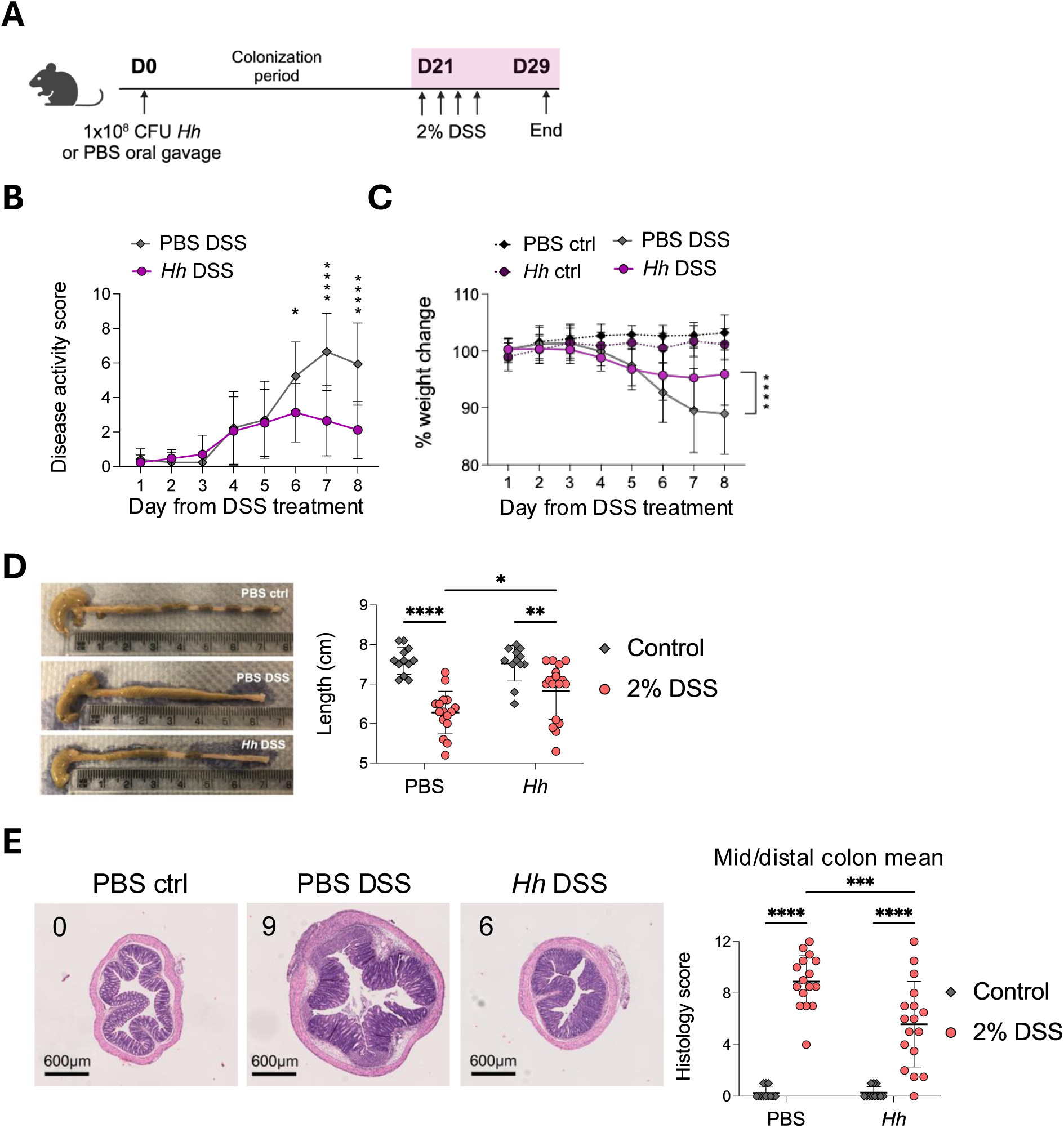
*H. hepaticus* attenuates DSS-induced inflammatory disease severity. C57BL/6 littermates were infected with 1x10^8^ CFU *H. hepaticus* or given PBS by oral gavage and left for 21 days to allow bacterial colonization. On day 21 mice were administered 2% DSS in drinking water or given normal water as controls. After 4 days mice were switched back to normal drinking water for a final 4 days before being culled on day 29. (A) Schematic of experimental design. (B and C) Following DSS treatment mice were assessed daily for clinical disease activity (B) and weight change (C). (D) Representative pictures of DSS-treated colons are shown (left) with quantification of length (right) taken at day 29. (E) Representative haematoxylin and eosin (H&E) sections of distal colon from uninfected control, uninfected DSS, and *Hh* DSS groups are shown (left). Histopathology of the mid and distal colon was assessed and presented as a mean score out of 12 (right). The histology scores of the representative H&E sections are highlighted in the top left of each image. Data are presented with mean ± SD and are pooled from 3 independent experiments with n=4-6 per experiment. Each data point represents an individual mouse (D and E) or data are shown as the daily mean of the experimental group (B and C). Statistical significance was determined by 2-way ANOVA (significance *p< 0.05, **p< 0.001, ***p< 0.001, ****p< .0001).

### *Hh* colonization reduces accumulation of inflammatory leukocytes in the colon after DSS treatment

To understand how *Hh* may be altering inflammatory responses at the cellular level, we undertook flow cytometric analysis of colonic lamina propria leukocytes (LPL) following DSS treatment. We designed flow cytometry panels to explore changes to myeloid cells and lymphocytes during DSS colitis and *Hh* infection (Supplementary Figure 1A-B). This approach allowed the identification of eosinophils, neutrophils, conventional dendritic cells (cDCs), monocytes and macrophages, B cells, CD4^+^/CD8^+^ T cells, and innate lymphoid cells (ILCs) (Figure 2A and Supplementary Figure 1A-B). We then performed dimensionality reduction analysis using t-distributed stochastic neighbour embedding (tSNE) to visualise cell populations (Figure 2A and Supplementary Figure 1C-D). In uninfected animals, DSS treatment caused marked increases in the proportions of neutrophils, monocytes, and intermediate monocyte-macrophages within the myeloid LPL compartment [26] (Figure 2A and 2B). In comparison, DSS treatment of *Hh* colonized mice resulted in significantly reduced proportional increases of these cell types (Figure 2A and 2B). Accordingly, within the DSS-treated groups, those colonized with *Hh* showed significantly reduced absolute numbers of colonic neutrophils and monocytes (Figure 2B).

**Figure 2:**
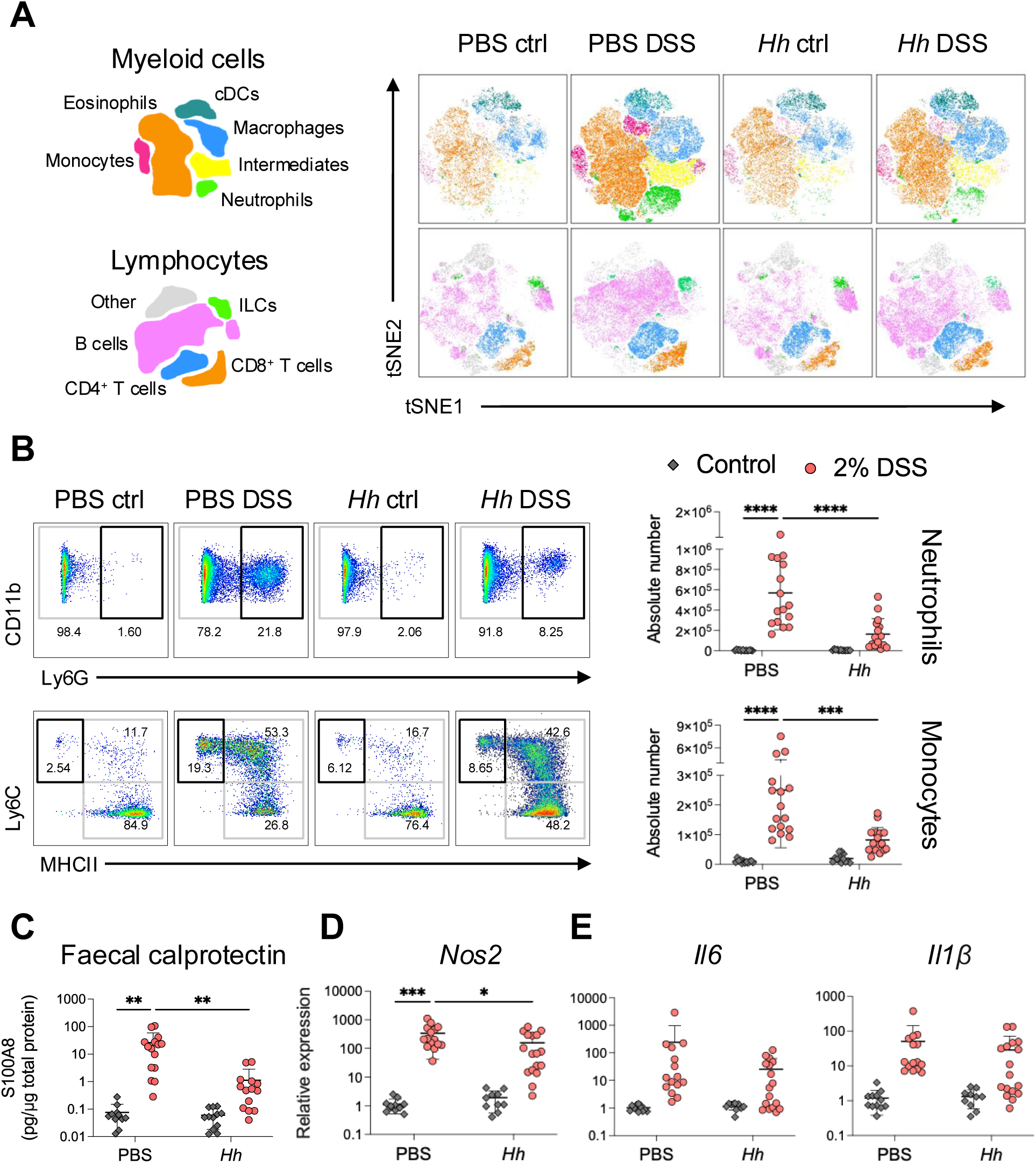
*H. hepaticus* colonized mice show reduced myeloid driven inflammatory responses. C57BL/6 littermates were infected with 1x108 CFU *H. hepaticus* or given PBS by oral gavage and administered 2% DSS in drinking water from days 21-24 or given normal water as controls. Mice were culled on day 29. (A and B) Flow cytometric analysis of CD45^+^ colonic LPL populations. (A) Schematics of myeloid and lymphoid populations analysed by tSNE are shown (left) with corresponding representative tSNE visualisation (right). (B) Representative dot plots of colonic CD11b^+^Ly6G^+^ neutrophils (upper) and CD11b^+^CD64^+^ monocyte-macrophages divided by expression of Ly6C and MHCII (lower) are shown (left) with quantification of neutrophil and Ly6C^+^MHCII^-^ monocyte frequency and absolute number (right). (C) Stool was collected at experiment endpoint and the concentration of S100A8 measured by ELISA. Concentrations are shown relative to total protein content determined by BCA assay. (D and E) A section of distal colon was isolated and expression of indicated genes measured using qPCR. Differences in gene expression were determined using the 2^-^ _ΔΔC(t)_ method, with gene expression normalised to the housekeeping gene *Rps29* and data shown as fold change relative to the PBS control group. Data are presented with mean ± SD and are pooled from 3 independent experiments with n=4-6 per experiment. Each data point represents an individual mouse. Statistical significance was determined by 2-way ANOVA (significance *p< 0.05, **p< 0.001, ***p< 0.001, ****p< .0001).

Numbers of eosinophils, cDCs, intermediate monocyte-macrophages, and mature macrophages were significantly increased in the LPL compartment of both uninfected and *Hh* colonized mice following DSS treatment (Supplementary Figure 2A). Of these cell types, eosinophils and intermediate monocyte-macrophages showed a significant reduction in DSS-induced expansion with *Hh* colonization, while macrophage and cDC numbers remained similar regardless of *Hh* colonization (Supplementary Figure 2A). Analysis of lymphocyte cell numbers revealed that only B cells showed a significant increase in numbers within the LPL following DSS treatment, and this was significantly lower in *Hh* colonized animals (Supplementary Figure 2A). Numbers of CD4^+^ and CD8^+^ T cells, as well as ILCs, were not significantly altered by either *Hh* colonization or DSS treatment (Supplementary Figure 2A). Overall, DSS treatment resulted in increased accumulation of various colonic immune populations, chiefly inflammatory myeloid cells and B cells. Many DSS-increased populations were significantly reduced with *Hh* colonization, with neutrophils and monocytes in particular showing a marked reduction in *Hh* colonized mice.

As DSS treatment caused a notable increase in colonic myeloid infiltration, we next assessed functional markers of myeloid driven inflammation. Faecal calprotectin is associated with neutrophilic inflammation and is widely used as a biomarker for IBD, as it correlates with endoscopic assessment of disease activity and is non-invasive [30, 31]. Calprotectin is a complex consisting of the S100A8 and S100A9 proteins, and we observed significantly increased concentrations of S100A8 in the stool of uninfected mice following DSS treatment (Figure 2C). However, *Hh* colonized mice showed significantly attenuated levels of faecal S100A8 after DSS (Figure 2C). We next measured colonic expression of *Nos2*, encoding inducible nitric oxide synthase (iNOS), an enzyme which is activated following exposure to inflammatory cytokines [32]. *Nos2* is upregulated in intestinal monocytes during acute inflammation [33] and we observed a clear upregulation of this gene in DSS-treated colons, although this was again significantly reduced in *Hh* colonized mice (Figure 2D). Finally, we measured colonic transcription of cytokines associated with myeloid inflammatory responses. Both *Il6* and *Il1β* showed a relative increase in transcription following DSS treatment, although this did not reach statistical significance (Figure 2E). While there were no significant changes to their expression with *Hh* colonization, there was a trend towards reduced upregulation of both cytokines after DSS treatment, with many *Hh* colonized mice showing comparable *Il6* and *Il1β* expression to non-DSS controls (Figure 2E). In contrast, colonic expression of *Tnfα*, a cytokine produced by monocytes and macrophages during colitis [17], was upregulated to a similar extent in both uninfected and *Hh*-colonized mice following DSS treatment (Supplementary Figure 2B). Taken together, these data indicate that *Hh* decreases the extent of myeloid infiltration to the colon during DSS colitis and reduces several markers of inflammation associated with neutrophils and monocytes.

### At steady state, *Hh* alters colonic myeloid cell dynamics and functions

We next investigated whether *Hh* drives changes to the colonic myeloid compartment at steady state, which could underlie the attenuated inflammatory responses observed during DSS colitis. We inoculated mice with *Hh* or PBS and examined the colonic myeloid pool at 21 days post-infection – the timepoint at which DSS was administered previously – while also performing the same analyses at day 7 to gain an understanding of the response kinetics over time. At day 7, we observed significant increases in the number of colonic neutrophils and intermediate monocyte-macrophages with *Hh* colonization, as well as a non-significant increase in monocyte numbers (Figure 3A). At 21 days post-infection, monocyte numbers remained elevated in *Hh*-infected mice, while neutrophils and intermediate monocyte-macrophages were no longer significantly increased compared to uninfected controls (Figure 3A). Despite *Hh* driving changes to numbers of colonic monocytes and intermediates, which are precursors of mature intestinal macrophages [34], we observed no changes to macrophage numbers at either timepoint (Figure 3A). Importantly, *Hh* colonization did not cause histological pathology at either timepoint (data not shown), indicating that this early infiltration of monocytes and neutrophils occurs in the absence of pathological inflammation. To further explore the response to steady state *Hh* colonization, we analysed colonic cytokine transcription using qPCR. We found that at day 7, *Hh* drove a significant upregulation of *Ifng* within the colon, while at day 21, transcription of *Tnfa* and *Il17a* were increased (Figure 3B). We then confirmed that colonization levels of *Hh* do not change significantly between day 7 and day 21 post-infection, indicating that changes to immune responses at these timepoints are not driven by bacterial load but instead likely represent priming of different immune populations over time (Figure 3C). Thus, *Hh* drives increased myeloid infiltration to the colon under homeostatic conditions, together with the induction of specific cytokine responses at different timepoints.

**Figure 3:**
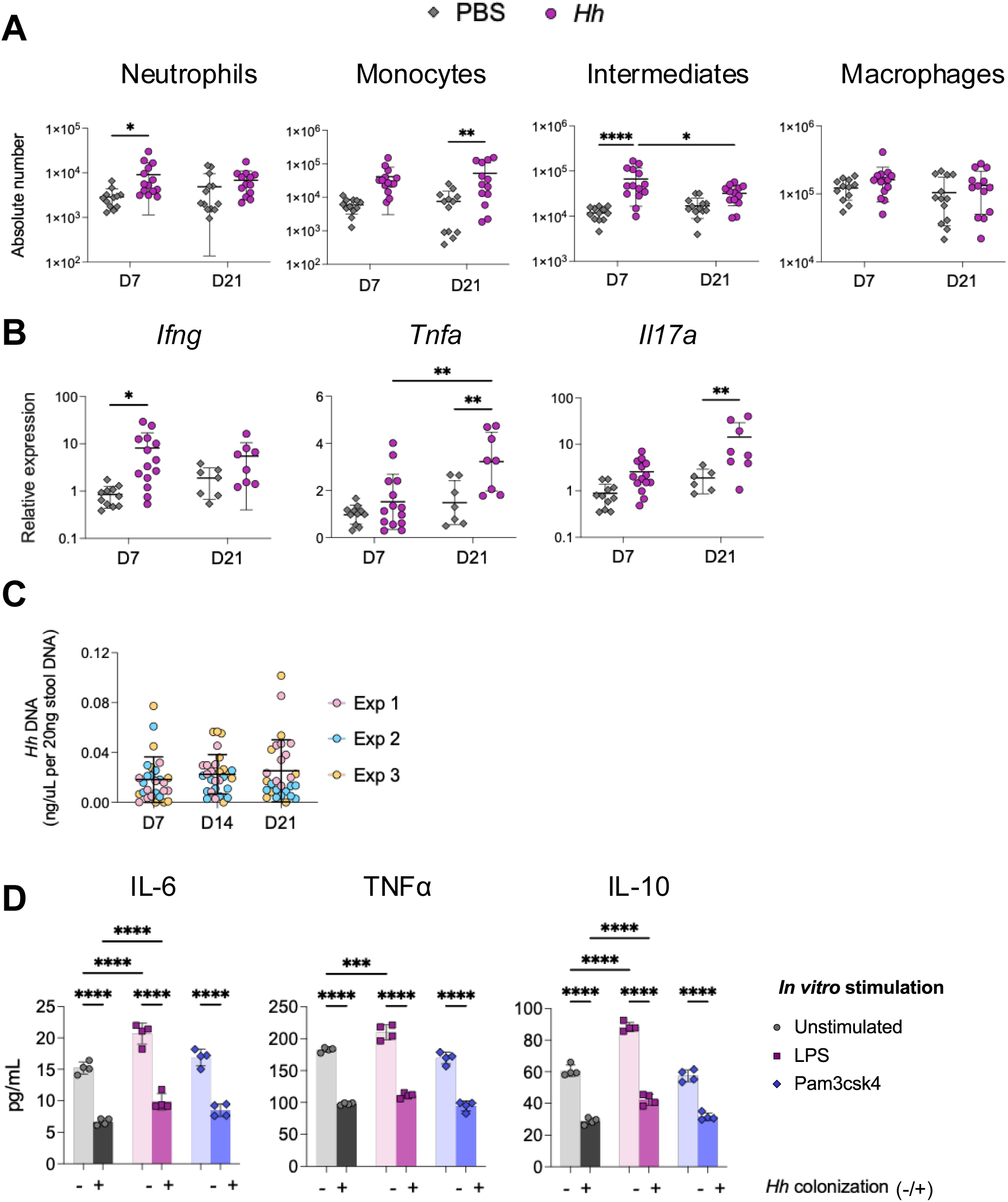
At steady state, *H. hepaticus* alters myeloid cell dynamics and functions. C57BL/6 littermates were infected with 1x10^8^ CFU *H. hepaticus* or given PBS by oral gavage and culled at day 7 or day 21 (A-D). (A) Absolute numbers of indicated colonic myeloid populations analysed by flow cytometry. (B) A section of distal colon was isolated and expression of indicated genes measured using qPCR. Differences in gene expression were determined using the 2^-^ _ΔΔC(t)_ method, with gene expression normalised to the housekeeping gene *Rps29* and data shown as fold change relative to the PBS control group. (C) *Hh* colonization was quantified in faecal DNA by qPCR of the *Hh*-specific *CdtB* gene. Individual experiments are shown in different colours. (D) 3x10^4^ FACS-isolated colonic macrophages (CD11b^+^CD64^+^Ly6C^-^MHCII^+^) were cultured for 16h either unstimulated or in the presence of 100ng/mL E. coli LPS or 100ng/mL Pam3csk4. The concentrations of indicated cytokines in culture supernatants were then measured by CBA. Macrophages were derived from PBS control or *Hh* colonized mice (*Hh* colonization indicated by +). Data are presented with mean ± SD and are pooled from 3 independent experiments with n=4-6 per experiment, each data point represents an individual mouse (A). Data are presented with mean ± SD and are representative of 2 independent experiments, data points are technical replicates pooled from 4 mice per group (E). Statistical significance was determined by 2-way ANOVA (significance *p< 0.05, **p< 0.001, ***p< 0.001, ****p< .0001) (A and E).

Finally, we tested whether *Hh*-driven changes to colonic myeloid infiltration were accompanied by altered functionality. During inflammation, increased colonic monocyte recruitment leads to the accumulation of macrophages with a pro-inflammatory phenotype, differing from that of mature macrophages present at steady state [35, 36]. We therefore asked whether the increased monocyte recruitment driven by steady state *Hh* colonization, in the absence of pathology, would invoke changes to differentiated macrophage functions. We infected mice with *Hh* and used fluorescence activated cell sorting (FACS) to isolate mature colonic CD11b^+^CD64^+^Ly6C^-^MHCII^+^ macrophages on day 21 post-infection. Macrophages were cultured overnight in medium alone or treated with different pathogen associated molecular patterns (PAMPs), including LPS [37] and Pam3csk4 [38], and cytokine secretion assayed. Across all treatment groups, we found that colonic macrophages from *Hh* colonized mice showed significantly reduced cytokine secretion compared to those from uninfected counterparts (Figure 3D). Interestingly, this appeared to be independent of any stimulus, as we observed this difference even in unstimulated cells (Figure 3D). Despite this, macrophages from both groups of mice produced significantly increased IL-6 and IL-10 following LPS stimulation, while the responses to Pam3csk4 were similar to those of unstimulated cells (Figure 3D). TNFα levels remained statistically similar between unstimulated and LPS-treated macrophages from *Hh* colonized donors, while those from uninfected mice produced significantly increased levels of this cytokine with LPS treatment (Figure 3D). These results indicate that *Hh* colonization reduces the ability of colonic macrophages to produce both inflammatory cytokines and IL-10 upon PAMP stimulation.

### The ability of *Hh* to attenuate DSS colitis severity is not dependent on TLR2

Having determined that *Hh* colonized mice show altered macrophage functions, we hypothesised that *Hh* may be having a direct effect on these cells through a specific microbial sensing pathway, potentially causing functional differences which result in attenuated inflammatory responses. Indeed, it was reported that *Hh* produces a polysaccharide which is recognised by TLR2 and promotes an anti-inflammatory macrophage phenotype [18]. We therefore postulated that this pathway may be responsible for alterations to macrophage phenotypes and that this could contribute to reduced DSS disease severity in these mice. Thus, we assessed whether the TLR2 pathway was required to drive the attenuation of DSS colitis by *Hh*.

To inhibit TLR2 signalling in this context, we utilised an anti-TLR2 monoclonal antibody (mAb) to block TLR2 signals *in vivo* [39]. We first validated the efficacy of αTLR2 mAb treatment by challenging mice with the specific TLR2 agonist Pam3csk4 [38] and analysing serum cytokine responses (Supplementary Figure 3A). While Pam3csk4 treatment drove increased levels of IL-6 and MCP-1 in the serum of control mice, these responses were almost completely ablated in mice pre-treated with αTLR2 mAb (Supplementary Figure 3A), confirming that this strategy could be employed to block TLR2 signals *in vivo*.

Mice were infected with *Hh* or given PBS as previously before all animals were challenged with 2% DSS from days 21-24 and culled on day 29. For the entire experiment duration, mice either received weekly doses of αTLR2 mAb or received an isotype control mAb (Figure 4A), and we confirmed that TLR2 blockade did not affect levels of *Hh* colonization (Supplementary Figure 3B). As expected, mice infected with *Hh* and given isotype control mAb showed significantly reduced disease activity and weight loss following DSS treatment compared to uninfected animals (Figure 4B). Treatment of uninfected mice with αTLR2 mAb had no significant impact on the severity of DSS symptoms, although there was a trend towards reduced disease activity and weight loss and a similar trend was present in the *Hh* infected αTLR2 mAb treated group (Figure 4B). Nevertheless, TLR2 blockade did not abrogate the protective effect of *Hh* colonization, as *Hh*-infected mice which received αTLR2 mAb showed significantly reduced DSS disease compared to uninfected αTLR2 mAb treated mice (Figure 4B). Furthermore, in the isotype control groups, mice colonized with *Hh* showed increased colon length compared to uninfected mice following DSS treatment (Figure 4C). Uninfected mice treated with αTLR2 mAb exhibited slightly increased colon length, while *Hh* colonized mice subjected to TLR2 blockade had comparable colon lengths to the *Hh*-infected isotype mAb treated group (Figure 4C). We then assessed the mid and distal colon for DSS-induced intestinal pathology. In the uninfected mice, we observed that treatment with αTLR2 mAb slightly reduced the severity of intestinal pathology, although not to a statistically significant level (Figure 4D). However, colonization with *Hh* significantly reduced the levels of histological inflammation, both in the isotype mAb and αTLR2 mAb-treated groups (Figure 4D). In parallel, we assessed the colonic lamina propria (cLP) for immune cell populations. *Hh* colonization significantly reduced the level of colonic neutrophils in both the isotype and αTLR2 mAb treated groups, while monocyte levels were reduced by both *Hh* colonization and αTLR2 mAb treatment (Figure 4E). Indeed, *Hh* colonized, αTLR2 mAb treated mice showed the lowest level of monocyte accumulation, suggesting an additive effect of *Hh* colonization and αTLR2 mAb treatment (Figure 4E).

**Figure 4:**
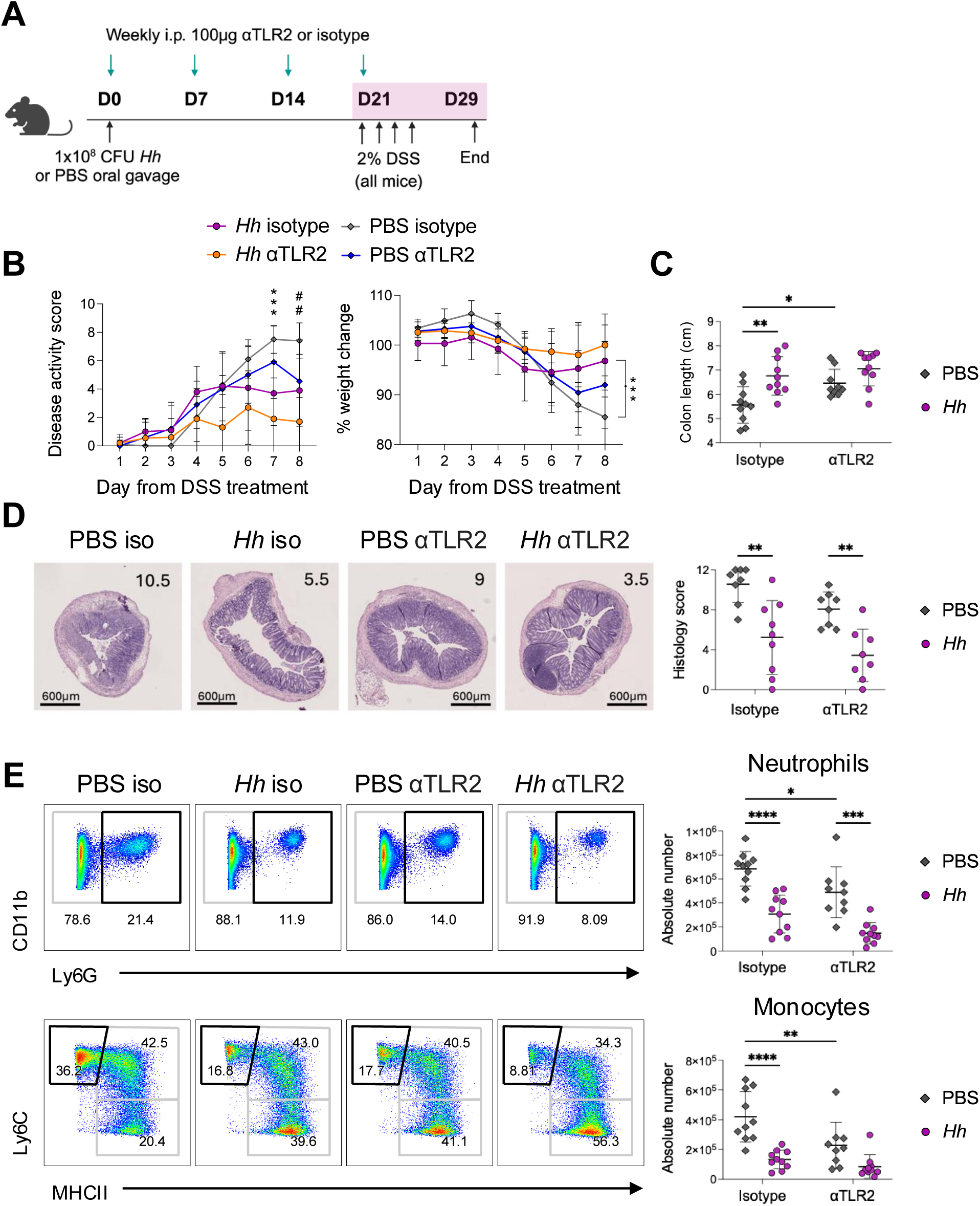
The ability of *H. hepaticus* to attenuate DSS severity is not dependent on TLR2. C57BL/6 littermates were infected with 1x10^8^ CFU *H. hepaticus* or given PBS by oral gavage. All groups were administered 2% DSS in drinking water from days 21-24 before being culled on day 29. Mice received either 100μg αTLR2 mAb or mouse IgG1, κ isotype control mAb by i.p. injection once weekly on days 0, 7, 14 and 21. (A) Schematic of experimental design. (B) *Hh* colonization was quantified in caecal content by qPCR of *CdtB*. (C) Following DSS treatment mice were assessed daily for clinical disease activity (left) and weight change (right). (D) Quantification of colon length. (E) Representative H&E sections of distal colon from PBS isotype, *Hh* isotype, PBS αTLR2, and *Hh* αTLR2 groups are shown (left). Histopathology of the mid and distal colon was assessed and presented as a mean score out of 12 (right). The histology scores of the representative H&E sections are highlighted in the top right of each image. (F) Representative dot plots of colonic CD11b^+^Ly6G^+^ neutrophils (upper) and CD11b^+^CD64^+^ monocyte-macrophages divided by expression of Ly6C and MHCII (lower) are shown (left) with quantification of neutrophil and Ly6C^+^MHCII^-^ monocyte absolute numbers (right). Data are presented with mean ± SD and are pooled from 2 independent experiments with n=5 per experiment. Data are shown as the daily mean of the experimental group (C), otherwise each data point represents an individual mouse. Statistical significance was determined by 2-way ANOVA (significance *p< 0.05, **p< 0.001, ***p< 0.001, ****p< .0001).

Together, these data indicate that although TLR2 blockade may have a slight attenuating effect on DSS-induced intestinal pathology, this effect is not as significant as the attenuating effect of *Hh* colonization. Importantly, loss of TLR2 signalling did not inhibit the ability of *Hh* to protect against DSS induced disease severity, indicating that attenuation of inflammatory responses by *Hh* in this context is not dependent on the TLR2 pathway.

### Persistent colonization by *Hh* is required for attenuation of DSS colitis and correlates with expansion of *Hh*-specific CD4^+^ T cells

We next considered whether the inhibitory effects of *Hh* colonization on myeloid responses might be driven indirectly through another cell type. As *Hh* colonization has been reported to induce a range of both regulatory and effector CD4^+^ T cell responses during homeostasis and inflammation [14, 15, 40], we assessed these in the context of DSS colitis. Although we did not observe changes to the total number of colonic CD4^+^ T cells following *Hh* infection with or without DSS treatment (Supplementary Figure 2A), we postulated that changes to CD4^+^ T cell phenotypes might alter the inflammatory responses driven by DSS. First, we used a major histocompatibility complex (MHC) class II tetramer containing the *Hh* peptide HH1713 to identify and characterise *Hh-*specific tetramer^+^ CD4^+^ T cells in the cLP and colon draining mesenteric lymph node (cMLN) during DSS colitis (Figure 5A). Although we observed comparable numbers of *Hh*-specific tetramer^+^ CD4^+^ T cells in the cLP, we found significantly increased numbers of *Hh*-specific tetramer^+^ CD4^+^ T cells in the cMLN during DSS colitis (Figure 5A), suggesting there may be increased exposure to *Hh* antigens following epithelial barrier damage. It has previously been described that the majority of *Hh-*specific colonic T cells differentiate into FOXP3^+^ RORγt^+^ Tregs at steady state, and that this phenotype does not change during DSS colitis [15]. In our hands, as reported, most HH1713 tetramer^+^ CD4^+^ T cells were FOXP3^+^ RORγt^+^ in both the cLP and cMLN, and DSS treatment did not significantly change the proportion of HH1713 tetramer^+^ cells expressing Foxp3 or RORγt (Supplementary Figure 4A).

**Figure 5:**
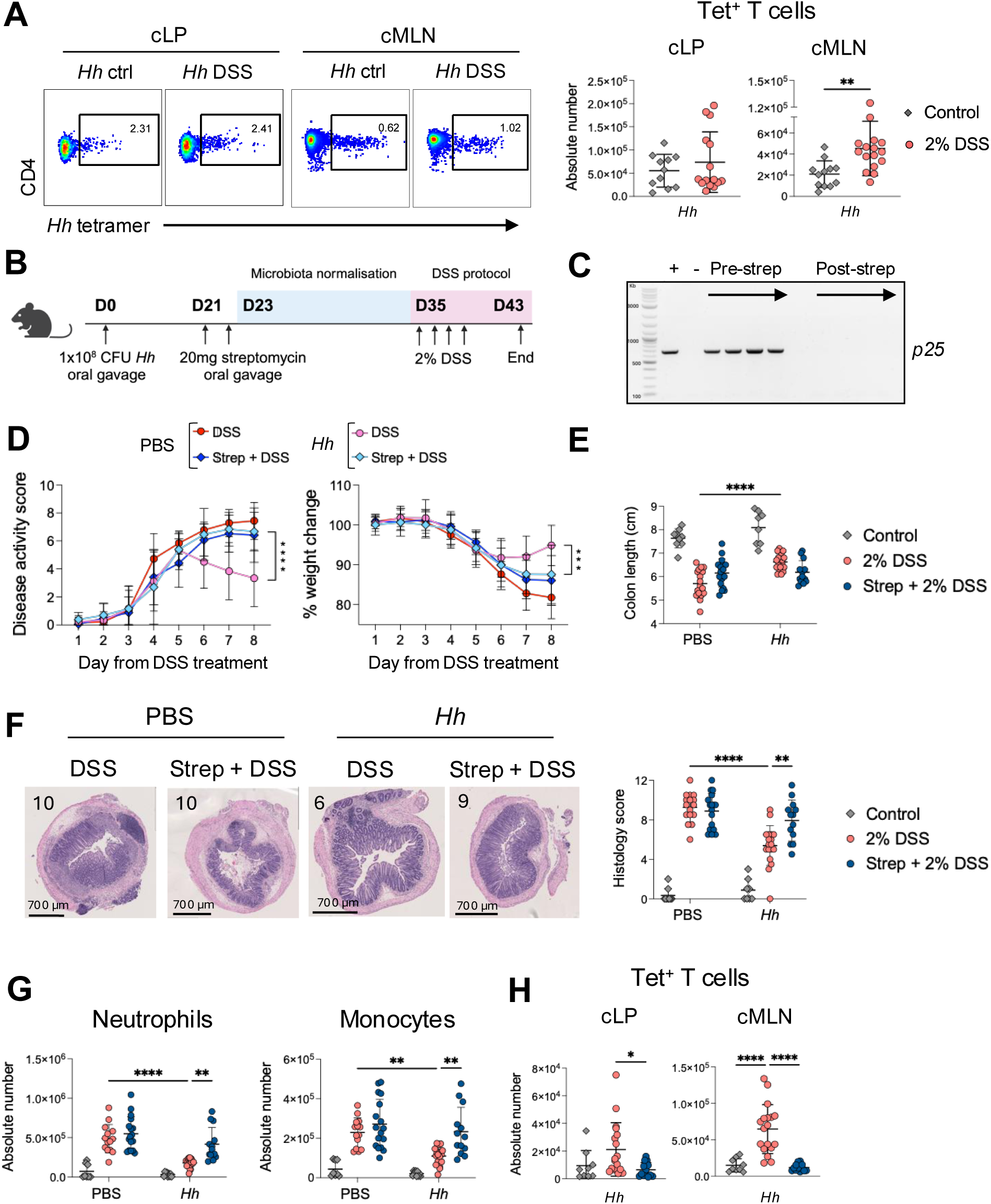
Persistent colonization by *Hh* is required for attenuation of DSS colitis and correlates with expansion of *Hh*-specific CD4^+^ T cells. (A) C57BL/6 littermates were infected with 1x10^8^ CFU *Hh* and administered 2% DSS in drinking water from days 21-24 or given normal water as controls. Mice were culled on day 29. *Hh* specific CD4^+^ T cells were identified using a fluorescently conjugated HH1713 tetramer. Representative dot plots of colonic LP and cMLN HH1713 tetramer^+^ (Tet^+^) CD4^+^ T cells are shown (left) with quantification of absolute numbers (right). (B – G) C57BL/6 littermates were infected with 1x10^8^ CFU *Hh* or given PBS by oral gavage, before certain groups received 20mg streptomycin by i.p. injection on days 21 and 22. Streptomycin treated mice were then co-housed with an experimentally naïve littermate for 2 weeks to promote microbiota normalization following antibiotic treatment. Naïve littermates were then removed and experimental mice administered 2% DSS in drinking water from days 35-38 or given normal water as controls. Mice were culled on day 43. (B) Schematic of experimental design. (C) *Hh* colonization status was assessed by PCR of the *Hh*-specific *p25* gene in faecal DNA, a representative image is shown. (D) Following DSS treatment mice were assessed daily for clinical disease activity (left) and weight change (right). (E) Quantification of colon length. (F) Representative H&E sections of distal colon from PBS DSS, PBS streptomycin + DSS, *Hh* DSS, and *Hh* streptomycin + DSS groups are shown (left). Histopathology of the mid and distal colon was assessed and presented as a mean score out of 12 (right). The histology scores of the representative H&E sections are highlighted in the top right of each image. (G) Absolute numbers of colonic LP CD11b^+^Ly6G^+^ neutrophils (left) and CD11b^+^CD64^+^Ly6C^+^MHCII^-^ monocytes (right) analysed by flow cytometry. (H) Absolute numbers of colonic LP (left) and cMLN (right) HH1713 tetramer^+^ (Tet^+^) CD4^+^ T cells analysed by flow cytometry. Data are presented with mean ± SD and are pooled from 3 independent experiments with n=3-6 per experiment. Data are shown as the daily mean of the experimental group (D), otherwise each data point represents an individual mouse. Statistical significance was determined by 2-way ANOVA (significance *p< 0.05, **p< 0.001, ***p< 0.001, ****p< .0001).

We next hypothesised that the survival of *Hh*-specific CD4^+^ T cells, and their expansion during DSS colitis, might reflect continued exposure to *Hh* antigens. To test this, we treated *Hh* colonized mice with the antibiotic streptomycin to clear the *Hh* colonization (Figure 5B). We carried out this treatment on day 21 post-*Hh* infection, thus allowing the same period of time as previously for *Hh* driven immune conditioning to occur. Following streptomycin administration, mice were co-housed with uninfected non-experimental littermates for 2 weeks to promote microbiota normalisation. Finally, the mice were challenged with 2% DSS from days 35-38, or given normal water as controls, before being culled on day 43 (Figure 5B). We first confirmed that streptomycin treatment had indeed eradicated *Hh* from the mice which were previously infected, using *p25* in faecal DNA as an indicator of *Hh* colonization [41] (Figure 5C). We then evaluated whether prior clearance of *Hh* with streptomycin altered the severity of DSS colitis. Control uninfected mice that were treated with streptomycin prior to DSS administration exhibited comparable disease activity scores and weight loss to those in the untreated DSS group (Figure 5D), indicating that this streptomycin treatment regimen had little impact on the course of DSS disease. As expected, *Hh* colonization attenuated the extent of DSS-induced clinical symptoms and weight loss (Figure 5D). However, *Hh* colonized mice which were treated with streptomycin did not show attenuated DSS disease symptoms, instead displaying similar disease activity levels to uninfected mice (Figure 5D). In addition, *Hh*-infected mice which received streptomycin showed significantly greater weight loss compared to the untreated, *Hh* colonized mice (Figure 5D). Although *Hh* colonization attenuated the shortening of colon length induced by DSS, we observed no significant differences in colon length with streptomycin treatment in either uninfected or *Hh* colonized mice (Figure 5E).

Upon assessment of intestinal pathology in the mid and distal colon, we found that streptomycin treatment of uninfected mice had no significant effect on histological inflammation (Figure 5F). *Hh* colonization resulted in significantly reduced histology scores compared to uninfected controls, as anticipated. However, streptomycin treatment ablated the attenuation of DSS-induced pathology driven by *Hh* colonization, with animals in this group displaying similar histology scores to the uninfected DSS groups (Figure 5F). Finally, we measured immune cell populations in the cLP and cMLN. Again, control uninfected mice that were treated with streptomycin prior to DSS administration harboured comparable numbers of colonic neutrophils and monocytes as those in the untreated DSS group (Figure 5G). As expected, *Hh* colonization significantly reduced the accumulation of colonic neutrophils and monocytes driven by DSS (Figure 5G). In contrast, *Hh*-colonized mice which received streptomycin showed significantly higher accumulation of neutrophils and monocytes in the cLP, which were comparable to those found in the uninfected DSS groups (Figure 5G).

To determine whether *Hh* infection clearance impacts the *Hh-*specific T cell response, we analysed CD4^+^ T cells in the cLP and cMLN. As previously, we observed a significant increase in numbers of HH1713 tetramer^+^ CD4^+^ T cells in the cMLN during DSS colitis and a trend towards an increase in the cLP, although this did not reach statistical significance (Figure 5H). Notably, in both the cLP and cMLN, clearance of *Hh* ablated the expansion of tetramer^+^ CD4^+^ T cells driven by DSS. Instead, mice treated with streptomycin showed comparable numbers of HH1713 tetramer^+^ CD4^+^ T cells to non-DSS controls (Figure 5H). These changes appear to be specific to HH1713 tetramer^+^ CD4^+^ T cells, as streptomycin treatment prior to DSS did not alter the total number of CD4^+^ T cells compared to DSS treatment alone, in either the cLP or cMLN (Supplementary Figure 5A). Overall, these results indicate that clearance of *Hh* with streptomycin prior to DSS administration abrogates the disease attenuating effect of *Hh* colonization, suggesting that the ability of *Hh* to alter inflammatory responses is dependent on continued interactions with host immune cells. Removal of *Hh* also prevents the expansion of *Hh*-specific CD4^+^ T cells during DSS colitis, suggesting that *Hh*-specific CD4^+^ T cell responses may be a key factor in the ability of this microbe to modulate host inflammatory responses after DSS administration.

Upon assessment of total CD4^+^ T cell transcription factor phenotypes, we observed an increase in numbers of Foxp3^-^RORγt^+^ Th17 cells in the cMLN during DSS colitis, which was greater in *Hh* colonized mice and not significantly altered by streptomycin treatment (Supplementary Figure 5B). However, in the cLP, numbers of Foxp3^-^ RORγt^+^ Th17 cells were largely comparable between all treatment groups (Supplementary Figure 5B). Numbers of both Foxp3^+^RORγt^-^ and Foxp3^+^RORγt^+^ Tregs were increased in the cMLN during DSS colitis to a comparable level in uninfected and *Hh* colonized mice and were not significantly changed by streptomycin treatment (Supplementary Figure 5C-D). Changes to Treg numbers were less pronounced in the cLP, although *Hh* colonized mice showed increased numbers of both Treg subsets during DSS colitis compared to uninfected mice (Supplementary Figure 5C-D). Among the Foxp3^-^ CD4^+^ T cells, numbers of RORγt^-^ Tbet^+^ Th1 cells were slightly increased in the cMLN following DSS, although not to a statistically significant level, and were not significantly altered by either *Hh* colonization or streptomycin treatment (Supplementary Figure 5E). In the cLP, the highest numbers of RORγt^-^Tbet^+^ cells were found in the streptomycin treated, *Hh* colonized mice, with all other groups showing comparable numbers of these cells regardless of *Hh* infection or DSS treatment (Supplementary Figure 5E). Finally, numbers of RORγt^+^Tbet^+^ cells, which are reported to produce both IL-17A and IFNγ and play a pathogenic role during colitis [14], were increased in the cMLN during DSS colitis and unaffected by streptomycin treatment in uninfected mice (Supplementary Figure 5F). However, in *Hh* colonized mice, numbers of these cells accumulated to a higher level during DSS colitis than in uninfected counterparts and were significantly reduced by streptomycin treatment (Supplementary Figure 5F). In contrast, in the cLP, numbers of RORγt^+^Tbet^+^ cells were comparable between infected and uninfected animals, and between control and DSS treated mice, with numbers of these cells only increasing following streptomycin treatment of *Hh* colonized mice (Supplementary Figure 5F). These results show that streptomycin treatment did not have a major impact on the dynamics of Treg or Th17 cell populations during DSS colitis, but increased the numbers of Tbet^+^ CD4^+^ T cells in the cLP of *Hh* colonized mice.

### Depletion of CD4^+^ T cells ablates the disease protective effect of *Hh* colonization

To test whether CD4^+^ T cell responses are necessary for *Hh* mediated protection, we employed an anti-CD4 mAb to deplete CD4^+^ T cells *in vivo* prior to DSS colitis [42]. Following three weeks of infection with *Hh*, mice received two doses of αCD4 mAb, given two days prior to and two days post induction of DSS treatment (Figure 6A). We used flow cytometry to confirm that the αCD4 mAb treatment had successfully depleted CD4^+^ T cells from the cLP (Figure 6B). While all mice in these experiments received DSS colitis, we observed that *Hh* colonization caused an increase in numbers of colonic CD4^+^ T cells among those which received the isotype control mAb treatment (Figure 6B).

**Figure 6:**
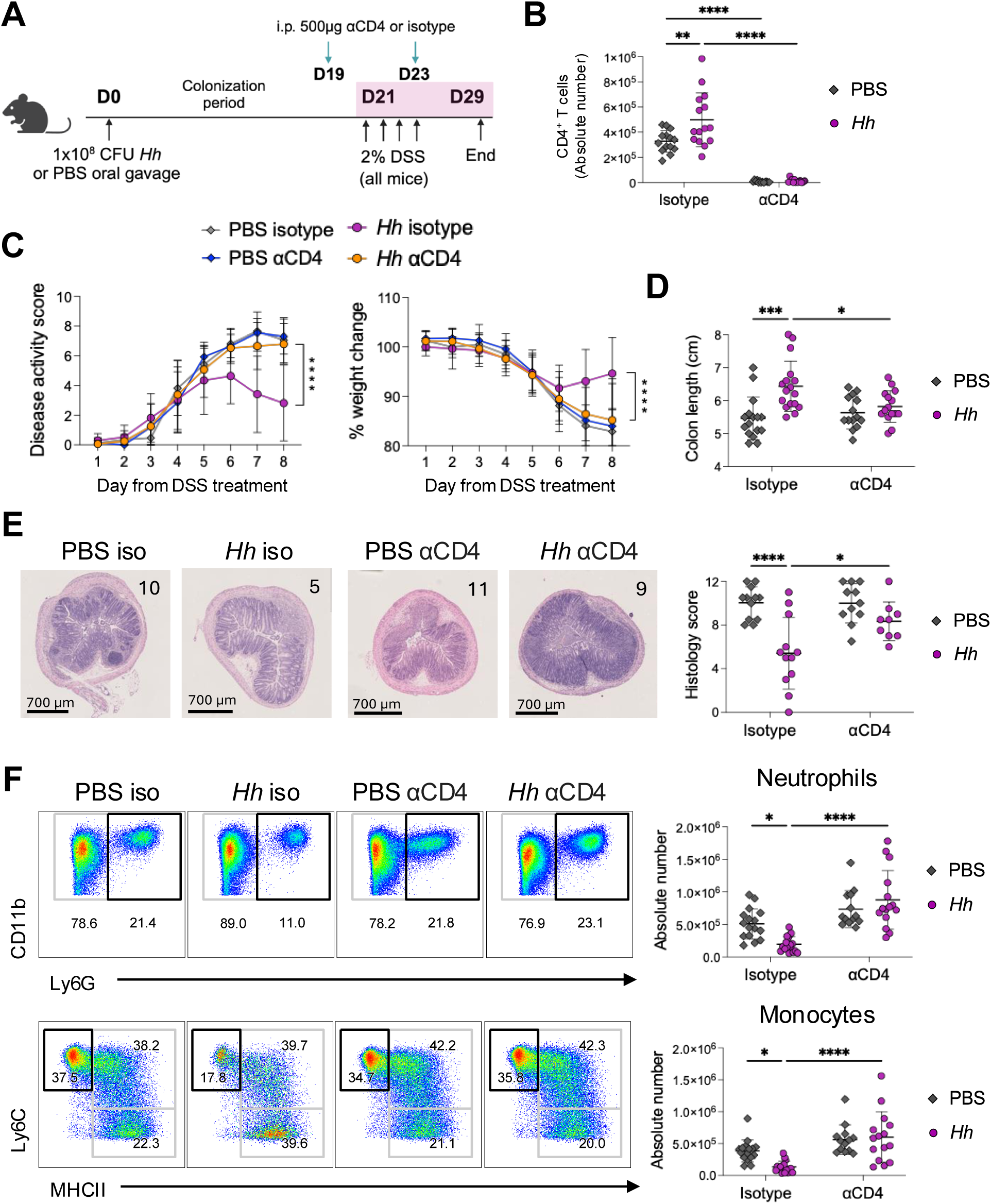
Depletion of CD4^+^ T cells ablates the disease protective effect of *Hh* colonization. C57BL/6 littermates were infected with 1x10^8^ CFU *H. hepaticus* or given PBS by oral gavage. All groups were administered 2% DSS in drinking water from day 21-24 before being culled on day 29. Mice received either 500μg αCD4 mAb or rat IgG2b, κ isotype control mAb by i.p. injection on days 19 and 23. (A) Schematic of experimental design. (B) Frequencies of colonic CD4^+^ T cells analysed by flow cytometry. (C) Following DSS treatment mice were assessed daily for clinical disease activity (left) and weight change (right). (D) Quantification of colon length. (E) Representative H&E sections of distal colon from PBS isotype, *Hh* isotype, PBS αCD4 and *Hh* αCD4 groups are shown (left). Histopathology of the mid and distal colon was assessed and presented as a mean score out of 12 (right). The histology scores of the representative H&E sections are highlighted in the top right of each image. (F) Representative dot plots of colonic CD11b^+^Ly6G^+^ neutrophils (upper) and CD11b^+^CD64^+^ monocyte-macrophages divided by expression of Ly6C and MHCII (lower) are shown (left) with quantification of neutrophil and Ly6C^+^MHCII^-^ monocyte absolute numbers (right). Data are presented with mean ± SD and are pooled from 3 independent experiments with n=5-8 per experiment. Data are shown as the daily mean of the experimental group (C), otherwise each data point represents an individual mouse. Statistical significance was determined by 2-way ANOVA (significance *p< 0.05, **p< 0.001, ***p< 0.001, ****p< .0001).

Upon DSS administration, we confirmed that *Hh* colonized mice that received the isotype control had significantly lower levels of disease activity and weight loss (Figure 6C). Among the uninfected mice, αCD4 mAb treatment had no effect on disease activity or DSS induced weight loss (Figure 6C). However, *Hh* colonized mice which received αCD4 mAb showed significantly greater DSS induced disease activity and weight loss compared to *Hh* infected control mice (Figure 6C). In fact, depletion of CD4^+^ T cells restored DSS disease activity and weight loss to a comparable level between *Hh*-infected and uninfected mice (Figure 6C). Similarly, colon lengths were greatest in the isotype treated *Hh* colonized group, while αCD4 mAb treatment of *Hh* infected mice caused a significant reduction in colon length which was comparable to the uninfected groups (Figure 6D). Furthermore, although αCD4 mAb treatment had no significant impact on histological inflammation in DSS-treated uninfected mice, the attenuated intestinal inflammation observed in *Hh* colonized mice was ablated by αCD4 mAb treatment (Figure 6E). Finally, we assessed numbers of neutrophils and monocytes in the cLP following DSS treatment. In the isotype control groups *Hh* colonization significantly reduced numbers of both cell types, but again αCD4 mAb treatment completely removed this effect (Figure 6F). Indeed, *Hh* colonized mice lacking CD4^+^ T cells showed comparable inflammatory infiltrates to uninfected mice following DSS (Figure 6F). Taken together, these results clearly demonstrate that CD4^+^ T cells are essential for the attenuation of DSS colitis induced by *Hh* colonization.

### At steady state, *Hh* drives Treg expansion and upregulates anti-inflammatory gene expression across effector T cell populations

To further explore colonic T cell responses during *Hh* colonization, and determine whether *Hh* drives specific functional changes which could underlie the attenuation of DSS colitis, we transcriptionally profiled T cells of mice colonized with *Hh* at steady state. Mice were infected with *Hh* or given PBS as previously, before we FACS-isolated CD3^+^ T cells from the cLP on day 21 and performed single cell RNA-sequencing (scRNA-seq). Unsupervised clustering of total CD3^+^ T cells revealed 18 transcriptionally distinct clusters (Figure 7A). Ten CD4^+^ clusters were identified, which broadly formed four major groups: Naïve T cells (cluster 5), T effector cells (clusters 3, 7, 11, 13, 14), proliferating T cells (cluster 12), and T regulatory cells (clusters 0, 6, 9) (Figure 7A).

**Figure 7:**
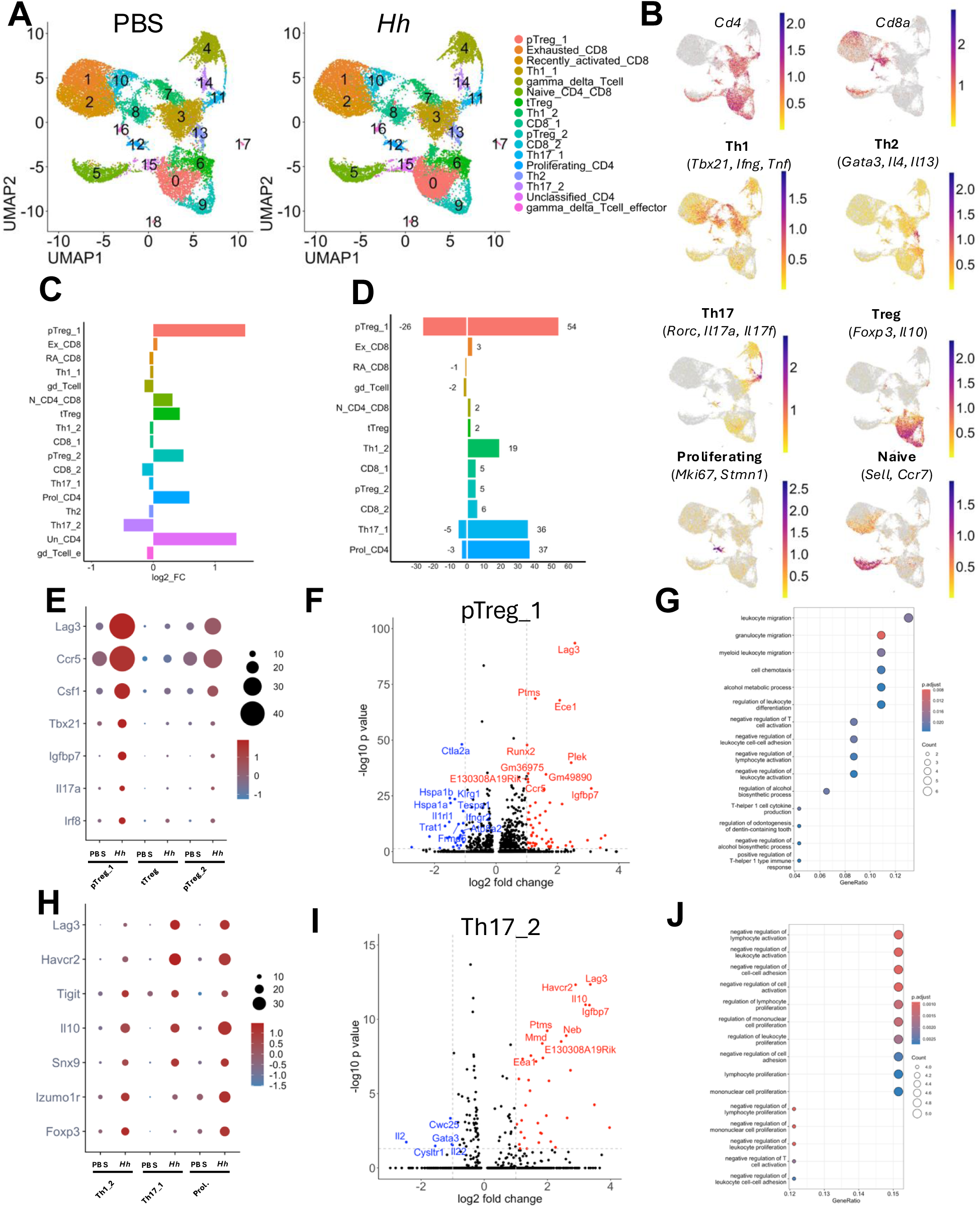
At steady state, *Hh* drives Treg expansion and upregulates anti-inflammatory gene expression across effector T cell populations. C57BL/6 littermates were infected with 1x10^8^ CFU *H. hepaticus* or given PBS by oral gavage and culled at day 21, total CD3^+^ T cells were sorted from cLP and underwent scRNA-sequencing. (A) UMAP visualization of total T cell subsets in cLP of PBS control (left) and *Hh* colonized (right) mice. (B) Feature plots of relative expression data overlayed on UMAP visualization of T cell subsets for selected markers. (C) Log2-transformed fold change of number of cells per cluster in *Hh* colonized compared with PBS control mice. (D) Number of differentially expressed genes per cluster in *Hh* colonized compared with PBS control mice. (E) Relative mean expression of selected markers in CD4^+^ T regulatory cell clusters. (F) Volcano plot of differentially expressed genes in cluster 0 (pTreg_1) cluster. (G) Pathway analysis of upregulated differentially expressed genes in cluster 0 (pTreg_1) cluster. (H) Relative mean expression of selected genes in T effector cell clusters. (I) Volcano plot of differentially expressed genes in cluster 11(Th17_2) in *Hh* colonized vs PBS control mice.

Regulatory T cells formed three clusters, two comprising peripherally induced Tregs (pTregs) based on *Rorc* expression (Clusters 0 and 9), and one thymically derived Treg (tTreg) (defined by *Ikzf2* (Helios) expression (cluster 6) (Figure 7A-B, Supplementary Figure 6B). The pTreg_1 cells (cluster 0) were the most enriched in *Hh* colonized mice (Figure 7C). Differential expression analysis between clusters from uninfected and *Hh* colonized mice revealed that pTreg_1 also had the largest number of differentially expressed genes (DEGs) (54 upregulated and 26 downregulated) (Figure 7D). The most significantly upregulated gene in pTreg_1 after *Hh* colonization was *Lag3* (Figure 7E-F), which acts as an inhibitory coreceptor in T cell activation and is important for Treg suppressive functions [43]. *Csf1*and *Ccr5* were also upregulated in pTreg_1 after *Hh* colonization (Figure 7E), as well as genes associated with T effector subsets such as *Tbx21, Il17a, Igfbp7* and *Irf8* [44, 45] (Figure 7E). After *Hh* colonization these same genes were also upregulated, albeit to a lesser extent, in the pTreg_2 population (cluster 9) but not in tTreg cells (cluster 6) (Figure 7E). Pathway analysis of DEGs between pTreg_1 in uninfected and *Hh* colonized mice demonstrated enrichment of pathways associated with migration of immune cells and negative regulation of immune cell activation (Figure 7G).

CD4^+^ effector T cells formed six clusters comprised of two Th1 clusters (cluster 3 and 7), two Th17 clusters (cluster 11 and 14), one Th2 cluster (cluster 13) and one cluster of proliferating effector T cells (cluster 12) (Figure 7A-B). Among these, one Th17 cluster (Th17_1, cluster 11), one Th1 cluster (Th1_2, cluster 7) and proliferating T cells (Prol., cluster 12) showed significant transcriptional differences between *Hh* colonized and uninfected mice (Figure 7H-I and Supplementary Figure 6D). After *Hh* colonization a similar gene expression program was upregulated in these effector subsets, characterized by genes encoding inhibitory molecules such as *Lag3*, *Havcr2* (Tim3) and *Tigit* (Figure 7H). In addition, genes traditionally associated with Treg cells, including *Il10*, *Foxp3* and *Izumo1r* (Folr4), were also upregulated across these effector clusters after *Hh* colonization (Figure 7H).

Pathway analysis of DEGs in cluster Th17_2 between uninfected and *Hh* colonized mice demonstrated enriched pathways associated with negative regulation of immune cell activation, proliferation and cell adhesion (Figure 7J). Together, these results suggest that *Hh* colonization drives an enhancement of regulatory T cell functions at steady state and may also skew some Th1 and Th17 effector cells towards a more anti-inflammatory phenotype.

## Discussion

*Hh* is recognised as an exemplar pathobiont species which drives IBD-like inflammation in the absence of immune regulation, however, in normal hosts, chronic infection is well tolerated without pathology thanks to the induction of avid regulatory responses [12]. While immune regulation is critical for maintaining a healthy relationship with the microbiota, the effects of *Hh*-induced regulatory responses on subsequent immune function remain poorly understood. Here, we investigated the impact of *Hh* colonization on inflammatory disease outcomes in the intestine. We found that mice infected with this pathobiont were protected against severe disease during DSS colitis, in a manner which was dependent on continued *Hh* colonization and required CD4^+^ T cells.

The host-microbiota relationship plays a critical role in health and disease, with recent research emphasising the constant interplay of interactions through which microbes help to shape immune function [1]. The extent of microbial impact on disease outcomes in the gut was recently highlighted using the DSS colitis model, in which a large number of C57BL/6 mice from the same SPF facility showed highly variable disease levels when treated with DSS [5]. Microbiota sequencing was employed to identify specific microbes whose relative abundance correlated with worsened or improved disease outcomes, and monocolonization of germ-free mice with these species was shown to replicate their effect on DSS disease [5]. Importantly, none of these animals were colonized with *Hh*, but this corroborates other studies describing how the microbiota and specific bacterial species can impact outcomes in DSS colitis [21, 46, 47]. Our results demonstrate that *Hh* acts as another member of the microbiota with the capacity to alter DSS colitis severity. Acting as a disease attenuating factor in this context, this contrasts the association between *Hh* and colitis observed in immunodeficient mice [9, 10, 12].

The ability of *Hh* to impact pathology during DSS colitis is likely underpinned by modulation of the host immune compartment at steady state. DSS colitis is immune-mediated and driven primarily by infiltrating myeloid cells [26], and we observed reduced DSS-induced neutrophil and monocyte accumulation in *Hh* colonized mice compared to uninfected controls. Despite this, at steady state, we noted modest increases to colonic neutrophil and monocyte infiltration following *Hh* colonization, as well as changes to colonic proinflammatory cytokine transcription, including an early *Ifng* response which was later followed by elevated levels of *Tnfa* and *Il17a*. These data indicate that different pathways and cell types may be responding to *Hh* over time, with early innate-driven responses likely being followed by induction of specific B and T cell responses. Importantly, these responses were not accompanied by pathology, indicating that *Hh* drives homeostatic immune responses in the intestine [1]. Our findings with *Hh* are in accordance with reports that the microbiota drives steady state myelopoiesis [48-50], and stimulates monocyte recruitment to the colon in the absence of inflammation [34, 51].

We found that colonic macrophages from steady state *Hh* colonized mice showed suppressed *ex vivo* cytokine production compared to those from uninfected donors. This was in contrast to the increased global colonic cytokine transcription observed in these mice, suggesting that *Hh* may have distinct functional effects on different cell types. Nevertheless, Abolins *et al*. reported a similar effect driven by the microbiota, in which splenocytes from wild mice showed greatly reduced cytokine production when stimulated with PAMPs *ex vivo*, as compared to laboratory mice [25]. Similarly, another study employed a system in which laboratory mouse embryos are implanted into wild mice to produce laboratory strain mice with a wild mouse-like microbiota [23]. These mice were then challenged with influenza infection and those with the ‘wild-like’ microbiota had reduced levels of TNFα, IL-6, and other inflammatory mediators in the lung after infection compared to genetically identical mice with a standard SPF laboratory microbiota. This conferred a survival advantage over their SPF counterparts, which the authors postulate may be due to the reduction in hyper-active cytokine production [23]. Our data suggests that *Hh* colonization may have a limiting effect on the cytokine production of some cell types, such as macrophages, while simultaneously increasing steady state expression of inflammatory cytokines in others. Intestinal macrophages, unlike monocytes or macrophages from other tissues, are refractory to stimulation as a result of the large microbial burden in the gut [52, 53]. *Hh* may be one of the species in the microbiota which contributes to this effect, although, somewhat surprisingly, macrophage IL-10 production also appeared reduced. This cytokine has previously been shown to be upregulated in macrophages from *Hh* colonized mice at steady state [18], however, in that study the IL-10 production was measured at 5-days post *Hh* colonization, as opposed to the day 21 timepoint assessed here. Therefore, extended interactions with *Hh*, and priming of adaptive immune responses, may imprint a phenotype which is not present early after infection.

Several studies have examined the specific microbial sensing pathways which mediate responses to *Hh* in different contexts. TLR2 was initially reported as the key pattern recognition receptor (PRR) involved in sensing of *Hh* [54], although a later study demonstrated that TLR2 was not required for colitis driven by *Hh* colonization of susceptible mice [55]. A role for TLR2 has been described for maintaining intestinal homeostasis [56], so it is therefore possible that TLR2-mediated recognition of *Hh* does occur but preferentially drives a protective, rather than inflammatory, response. Consistent with this hypothesis, TLR2-mediated sensing of a *Hh* polysaccharide moiety has been reported to induce an anti-inflammatory phenotype in macrophages [18]. However, using a mAb to block TLR2 signalling *in vivo*, we demonstrate that the attenuation of DSS colitis by *Hh* is not dependent on TLR2 signalling. Nevertheless, even in the absence of *Hh* colonization, we observed a slight reduction in DSS disease severity in mice treated with αTLR2 mAb, which was most notable in uninfected animals. These results suggest that loss of TLR2 signalling has a slight alleviating effect on DSS disease, although much less prominent than the protective effect of *Hh* colonization. This is in contrast to some reports that *Tlr2*^-/-^ mice display increased mortality and disease severity when challenged with DSS [57, 58]. However, other studies describe similar findings to those shown here, with *Tlr2*^-/-^ and *Tlr4^-/-^* mice reported to show attenuated DSS disease compared to wild-type [59]. These discrepancies likely reflect different microbiota composition among the TLR knockout strains housed in different animal facilities, which plays a dominant role in susceptibility to DSS colitis [5, 21]. By using antibody-mediated blockade of TLR2 signalling in wild-type mice we were able to avoid this issue, and our results clearly demonstrate that the protective effect of *Hh* colonization on DSS colitis was independent of TLR2.

The induction of a diverse range of different CD4^+^ T cell responses by *Hh* has been well described [14, 15, 40], with Tregs being the dominant type of *Hh*-specific T cell in the cLP of healthy wild-type mice [15]. Together with *Hh*-driven changes to myeloid inflammatory responses and steady state macrophage phenotypes, we observed an expansion of *Hh*-specific CD4^+^ T cells following DSS treatment of *Hh* colonized mice. Using a MHCII tetramer system, we found no differences in expression of the key transcription factors RORγt or Foxp3 in *Hh*-specific CD4^+^ T cells during DSS colitis. Commensal-specific CD4^+^ Treg cells have previously been described to become ‘bystander activated’ during acute intestinal infection, resulting in acquisition of Tbet and RORγt expression and loss of Foxp3 [15, 60]. However, our results are consistent with a previous report that this phenotype switching does not occur in *Hh*- specific CD4^+^ T cells during DSS colitis [15]. The expansion of tetramer^+^ CD4^+^ T cells during DSS colitis as described here may instead be the result of increased exposure to *Hh*-derived antigens following DSS-induced barrier damage.

We found that clearance of *Hh* colonization by streptomycin treatment abrogated the disease protective effect during subsequent DSS colitis challenge, indicating that the disease attenuation requires continued host interactions with *Hh*. Importantly, although perturbations to the microbiota with antibiotics can worsen DSS colitis severity [61], we did not observe any differences in disease severity in uninfected mice treated with streptomycin prior to DSS. This may be due to the microbiota normalisation period we employed prior to DSS induction, in which streptomycin-treated animals were co-housed with uninfected littermates to restore their microbiota composition. However, clearance of *Hh* by streptomycin resulted in loss of protection from subsequent DSS challenge. This finding raises the hypothesis that persistent colonization by pathobionts like *Hh* is tolerated because the dynamic interactions between *Hh* and the host immune system establish a beneficial equilibrium in which the intestine becomes more resilient to other challenges or damage. The potential ‘cost’ of such an equilibrium is that defects or loss of immune regulatory pathways can elicit deleterious responses to pathobionts that drive chronic intestinal inflammation. The requirement for persistent *Hh* colonization for protection from DSS colitis also suggests that rather than long-term imprinting or training of local resident innate immune cells, there must be constant recognition and/or sensing of *Hh*-derived molecules to perpetuate this beneficial equilibrium. This led us to postulate that adaptive immune pathways, in particular the *Hh*-specific CD4^+^ T cell population that expanded during DSS colitis, could be a critical component of the protective mechanism.

Using an antibody to deplete CD4^+^ T cells *in vivo*, we demonstrate that these cells are essential for the attenuation of DSS colitis severity by *Hh*. Although depletion of CD4^+^ T cells in control uninfected mice had no significant impact on DSS disease, it completely abrogated the protective effects of *Hh* colonization observed across all clinical and inflammatory parameters. As noted above, we did not observe significant changes in canonical transcription factor expression in CD4^+^ T cells from *Hh*- colonized mice following DSS challenge. Therefore, to better understand how *Hh* colonization could functionally modulate intestinal CD4^+^ T cell responses, we conducted scRNA-seq analyses of cLP CD4^+^ T cells. Our data indicate that at steady state *Hh* drives the upregulation of an anti-inflammatory gene signature across a range of CD4^+^ T cell subsets, including Th1 and Th17 clusters, as well as peripherally induced regulatory T cell clusters. This *Hh* driven module is characterized by upregulation of genes associated with Tregs, including *Foxp3*, *Il10* and *Izumo1r* (Fr4) in Th1 and Th17 clusters. Furthermore, common across these clusters is upregulation of multiple genes encoding co-inhibitory receptors, including *Lag3*, *Havcr2* (Tim3) and *Tigit*. This is in line with a recent study which reported that small intestinal LP Th17 cells specific for segmented filamentous bacteria (SFB) have an anti-inflammatory phenotype characterised by expression of IL-10 and multiple co-inhibitory receptors [62]. This is further consistent with the idea that intestinal CD4^+^ T cells exhibit plastic phenotypes which are modulated by microbial responses, rather than as defined helper T cell subsets [63]. Our data suggest that *Hh* promotes an anti-inflammatory response across a broad range of CD4^+^ T cell populations, enhancing the regulatory T cell response in the cLP beyond that of canonical regulatory T cell populations. This modulation of the effector T cell compartment may be contributing to the persistence of *Hh* in the intestinal tract, while simultaneously providing a heightened level of quiescence to inflammatory insults, which in the case of colitis may be beneficial to the host. However, it will be important to characterise the T cell compartment of *Hh* colonized mice in more depth and determine whether these phenotypic changes alter responses to pathogenic infections where tolerance may not be advantageous. Regardless, this increase in immune regulation appears to then limit the extent of inflammation following DSS challenge and likely underpins the critical role for CD4^+^ T cells in *Hh* mediated disease attenuation.

Overall, our study provides evidence that colonization with the pathobiont *Hh* can provide a beneficial, disease attenuating effect for the host, by modulating inflammation during DSS colitis. We show that this protective effect requires continued colonization with *Hh* and is dependent on CD4^+^ T cells, which were found to exhibit anti-inflammatory changes to gene expression with *Hh* colonization. These data provide new insight into the effects of pathobiont species on immune functions and contribute to the understanding that the microbiota primes the intestinal immune system at steady state, thereby altering immune responses during inflammatory settings.

## Methods

### Mice

Wild-type C57BL/6 were obtained from Envigo (Huntingdon, UK) and maintained under specific pathogen free conditions at the Central Research Facility, University of Glasgow, UK. Procedures were performed under a Project License and Personal Licenses issues by the UK Home Office. Mice were routinely screened to confirm the absence of *Helicobacter spp.* Female mice were used for all experiments at age 6-12 weeks.

### *Hh* growth and infections

#### Bacterial culture

*Helicobacter hepaticus* type strain 51449 (ATCC) [7] was cultured in tryptone soya broth (Oxoid) containing 10% foetal bovine serum (FBS) and Skirrow Campylobacter selective supplement (Oxoid). Cultures were grown microaerophillically for up to 4 days at 37℃ in a shaking incubator at 110 RPM, as described [10].

#### Infection with Hh

The concentration of *H. hepaticus* cultures were estimated using the equation 1 OD_600_ = 1x10^8^ CFU/mL [64]. Bacteria were assessed for viability using the LIVE/DEAD BacLight Bacterial Viability kit (ThermoFisher) then centrifuged at 5000 G for 10 mins to pellet bacteria and resuspended at 5x10^8^ CFU/mL in sterile PBS. Mice were infected with 0.2mL *H. hepaticus* (∼1x10^8^ CFU) in PBS by oral gavage.

### Induction of colitis

Mice were given 2% dextran sulphate sodium (DSS) (molecular weight ∼40,000, Alfa Aesar) in drinking water for 4 days, with fresh 2% DSS supplied on day 2. On day 4, animals were switched back to normal drinking water until the experiment end point on day 8. Control animals received normal drinking water. Mice were weighed daily during the 8-day DSS protocol and assessed for clinical signs of DSS-induced disease, receiving a disease score out of 16. Disease scores were based on 4 parameters (weight loss, bleeding, stool consistency, general appearance) with a maximum score of 4 for each.

### Antibody treatments

Mice were treated with purified anti-mouse CD282 (TLR2) clone QA16A01 (mouse IgG1, κ) recombinant antibody (BioLegend) to block TLR2 signalling. Animals received 100μg antibody once weekly for the duration of the experiment (4 doses total). Control animals received purified mouse IgG1, κ isotype control antibody (BioLegend) at the same dose. To deplete CD4^+^ T cells, mice were treated with purified anti-mouse CD4 clone GK1.5 (rat IgG2b, κ) monoclonal antibody (BioXcell). Animals received 2 doses of antibody at 500μg, 2 days prior to and 2 days post commencement of DSS treatment. Control animals received purified rat IgG2b, κ isotype control antibody (BioXcell) at the same dose. All antibodies were administered by intraperitoneal injection in 0.2mL sterile PBS.

### *Hh* clearance by antibiotic treatment

Mice were treated with antibiotic Streptomycin sulphate salt (Sigma-Aldrich) for the purpose of clearing *H. hepaticus* infection. On day 21 following infection, mice received 2 doses of Streptomycin on consecutive days by oral gavage at 20mg per dose in 0.2mL PBS. Control animals received PBS. On day 23, treated animals were co-housed with non-treated littermates to help normalise the microbiota following antibiotic depletion. Mice were co-housed for 2 weeks before untreated littermates were removed and the DSS protocol commenced. Mice were screened prior to DSS treatment to confirm the absence of *H. hepaticus* in antibiotic-treated animals.

### Isolation of lamina propria leukocytes

Colons were harvested and opened longitudinally before being washed in PBS and cut into pieces. Tissues were stored at 4℃ in HBSS (no calcium, no magnesium) (ThermoFisher) supplemented with 10% FBS. For lamina propria leukocyte (LPL) isolation, tissues were transferred to HBSS supplemented with 2mM EDTA (ThermoFisher) and incubated at 37℃ in a shaking incubator at 220 RPM for 15 mins before repeating this step a 2^nd^ time in new EDTA-HBSS. Colon tissue was then digested at 37℃ for 15 mins in a shaking incubator at 220 RPM with 0.65mg/mL collagenase D, 0.5mg/mL collagenase V, 30μg/mL DNAse I, and 1mg/mL dispase (all Sigma-Aldrich). Digestion was performed in complete R10 medium (RPMI-1640 supplemented with 10% FBS, 100U/mL penicillin, 100μg/mL streptomycin and 2mM L-glutamine – all Gibco). Digested samples were passed through 100μm and 40μm filters to obtain single cell suspensions. Cells were then centrifuged twice in R10 at 400G for 10 mins at 4℃.

### Flow cytometry

Up to 5x10^6^ cells per sample were stained with surface antibodies for flow cytometry analysis. All staining steps were carried out in PBS supplemented with 2% FBS and 2mM EDTA (FACS buffer). Cells were first incubated with Fixable Viability Dye (Invitrogen) and TruStain FcX (anti-mouse CD16/32) antibody (BioLegend) for 20 mins at 4℃ to exclude dead cells and minimise non-specific antibody binding, respectively. Cells were washed at 400 G for 5 mins at 4℃ then stained with a combination of surface antibodies for 30 mins at 4℃. Cells were washed again and fixed in Fixation Buffer (BioLegend) for 30 mins at 4℃, then washed and stored in FACS buffer at 4℃ until acquisition. For intracellular staining of nuclear transcription factors, cells were fixed using the Foxp3/Transcription Factor Staining Buffer Set (eBioscience) for 1 hour at 4℃. Cells were washed and stained with intracellular antibodies resuspended in Permeabilization Buffer (eBioscience) for 1 hour at room temperature (RT), before being washed and resuspended in FACS buffer before acquisition. CD4^+^ T cells specific for *H. hepaticus* epitope HH1713 were measured by flow cytometry using an MHCII tetramer (NIH tetramer core). Isolated LPL were incubated with fluorescently labelled I-A^(b)^/HH1713 tetramer at 37℃ for 2 hours in R10 containing TruStain FcX at 1:100 (BioLegend), before staining with surface and intracellular antibodies as described above. All samples were acquired on a BD LSR Fortessa and analysed using FlowJo (version 10, BD Bioscience).

### Colonic macrophage sorting and stimulation

Colonic LPL were isolated and stained with Viability Dye, TruStain FcX, and surface antibodies as above. CD11b^+^CD64^+^Ly6C^-^MHCII^+^ macrophages were sorted using a BD FACS Aria IIU or FACS Aria III (BD). Macrophages were sorted into R10 and stored at 4℃ prior to counting and stimulation. FACS-isolated LPL macrophages were seeded at 3x10^4^ cells per well in 0.2mL R10. Macrophages were either left unstimulated, treated with 100ng/mL LPS from *Escherichia coli* (Sigma-Aldrich), or 100ng/mL Pam3csk4 (InvivoGen) for 16h at 37℃ with 5% CO_2_. Culture supernatants were harvested and stored at -20℃ for downstream analyses.

### Cytokine measurement

The concentrations of IL-6, TNFα, and IL-10 in stimulated macrophage culture supernatants were measured using the Cytometric Bead Array Flex Set kits (BD Biosciences) according to the manufacturer’s instructions. Supernatants were assayed undiluted, and data were acquired using a BD FACS Canto II.

### RNA extraction and cDNA synthesis

1cm sections of distal colon were harvested and stored in RNA*later* Stabilization Solution (Invitrogen) at -80℃. Tissue samples were transferred to QIAzol Lysis Reagent and homogenised in a TissueLyser twice for 1 min at 25Hz using a 5mm Stainless Steel Bead (all Qiagen). Samples were centrifuged at 12,000 G for 5 mins at RT and supernatants transferred to a new 1.5mL Eppendorf. Supernatants were mixed with chloroform and incubated for 3 mins at RT before centrifuging at 12,000 G for 15 mins at RT. The aqueous layer containing RNA was harvested and mixed with 100% ethanol at 1.5X volume. Purification of RNA was carried out using the RNeasy Mini Kit with DNA digestion using the RNase-free DNase Set (both Qiagen). RNA concentration and quality were measured using a NanoDrop Spectrophotometer. RNA was stored at -80℃ until use. cDNA was synthesised from RNA using the High-Capacity cDNA Reverse Transcription Kit (Applied Biosystems). 2000ng RNA was reverse transcribed and resultant cDNA was diluted 1/10 in nuclease-free H_2_0 and stored at -20℃.

### Detection and quantification of *Hh* DNA in caecal contents and stool

Caecal contents and stool were collected in 1mL PBS and stored at -20℃ until extraction. Caecal/faecal DNA was extracted and purified using the QIAamp PowerFecal Pro DNA Kit (Qiagen) according to manufacturer’s instructions. DNA concentrations were measured using a NanoDrop Spectrophotometer. *Hh* DNA was detected in stool DNA samples by PCR of the *p25* gene using forward primer ATGGGTAAGAAAATAGCAAAAAGATTGCAA, reverse primer CTATTTCATATCCATAAGCTCTTGAGAATC [41], and GoTaq Green Master Mix 2X (Promega). Amplicons were analysed by 1% agarose gel electrophoresis containing 0.01% SYBR safe DNA Gel Stain (Invitrogen). *Hh* DNA was quantified in caecal DNA samples by qPCR of the gene *CdtB* using forward primer TCGTCCAAAATGCACAGGTG and reverse primer CCGCAAATTGCAGCAATACTT [65]. To generate a standard curve, a serial dilution of *Hh* DNA was produced using DNA extracted from *Hh* bacterial cultures using the PureLink Genomic DNA Mini Kit (Invitrogen) following the manufacturer’s protocol for gram negative bacteria. A top concentration of 25ng/μL *Hh* DNA was used. Caecal DNA samples were added for a total of 20ng DNA per reaction. All primers were obtained from IDT.

### Quantitative RT-PCR

Quantitative real-time PCR (qPCR) was performed using PowerUp SYBR Green Master Mix and the QuantStudio 7 Flex Real-Time PCR System (both Applied Biosystems). Per sample the qPCR reaction mixture for each gene of interest contained: 5μL SYBR Green Master Mix 2X, 0.5μL forward and 0.5μL reverse primer at 10μM working concentration. cDNA samples were added at 4μL per reaction and analysed in triplicate, with gene expression levels for each sample normalised to the housekeeping gene *Rps29* (encoding ribosomal protein S29). Differences in gene expression were determined using the 2^-ΔΔC(t)^ method [66]. All primers were obtained from IDT.

### Protein quantification from faecal samples

#### Stool processing

Faeces were collected from inside the colon and stored in 1mL sterile PBS at -80℃ until processing. Stool samples were homogenised in PBS using a TissueLyser twice for 1 min at 25Hz using a 5mm Stainless Steel Bead (both Qiagen). Stool samples were centrifuged at 12,000 G for 15 mins at 4℃ and supernatants were transferred to a new 1.5mL Eppendorf and stored at -80℃ for subsequent protein quantification.

#### S100A8 ELISA

The concentration of S100A8 (Calprotectin subunit) in faecal samples was measured using the mouse S100A8 DuoSet ELISA kit (BioTechne) according to the manufacturer’s instructions. Samples from non-DSS controls were assayed undiluted, while samples from DSS-treated mice were assayed between 1/10 and 1/1000 dilution. Absorbance was measured at 450nm/540nm using a plate reader. S100A8 protein concentrations were normalised to the total protein content per sample, as measured by Pierce bicinchoninic acid (BCA) assay (Thermo Scientific) performed in parallel using the manufacturer’s protocol for assaying in microplates. BCA absorbance was measured at 562nm using a plate reader.

### Histology

#### Processing, embedding, and sectioning

Approximately 1cm sections of proximal, mid, and distal colon were cut using a scalpel blade. Fat and stool were gently removed and samples were transferred to 10% neutral buffered formalin (Sigma-Aldrich) and fixed for 24h at RT, before being transferred to 70% ethanol for storage at RT until processing. Tissue was embedded into paraffin blocks and sectioned at 5μm thickness using a microtome. Sections were loaded onto positively charged slides and dried overnight at RT before being stored at 4℃ until staining.

#### H&E staining

Slides were incubated in an oven at 60℃ for 30-60 mins to melt paraffin, before being deparaffinised and rehydrated through a series of incubations in xylene and a graded alcohol series (xylene 3 mins x2, 100% ethanol 3 mins x2, 90% ethanol 3 mins x2, 70% ethanol 3 mins x2, distilled water 2 mins). Slides were stained with Harris Haematoxylin for 2 mins before being rinsed in running water. Differentiation was carried out by incubating slides in 1% acid/alcohol, running water, and Scott’s Tap Water Substitute for 30 secs each. Slides were then counter stained by first incubating in 70% ethanol and then in Eosin Y Stain 1% (alcoholic) for 30 secs and 3 mins, respectively. Slides were then dehydrated using the reverse graded alcohol series described above until incubation in xylene. Coverslips were mounted over tissue sections using DPX Mounting Medium. All reagents were obtained from CellPath.

#### Histological colitis scoring

Following H&E staining, slides were digitized using a NanoZoomer slide scanner (Hamamatsu) and sections were viewed using Aperio ImageScope software (Leica Biosystems). Colon histopathology was assessed in a blinded, semiquantitative fashion as described [5]. Scores from the mid and distal colon were averaged to provide a mean histology score between 0 and 12 for each mouse, with 12 being the most severe and 0 showing no pathology.

### Single-cell RNA sequencing

Colonic LPL were isolated as described and sorted using FACS for CD3^+^ T cells. Each sample was pooled from 4 individual mice, for a total of 2 T cell samples from *Hh*-colonized mice and 2 from PBS control mice. Samples underwent library preparation by the NGS & Bioinformatics facility, University of Glasgow.

#### Pre-processing of 10x Genomics scRNA-seq

Single cell libraries, averaging 10k cells per sample, were prepared using the 10X Chromium Next GEM X Single Cell 3’ kit v4, sequenced on an Illumina NextSeq 2000 with read length 28x90bp, and to an average depth of 40k reads per cell. The fastQ files were aligned using Cell Ranger (V8.0.1), under default settings.

### Analysis of single-cell RNA sequencing data

#### Quality control and processing of scRNA-seq data

The filtered count data for each sample was imported into R (V4.5.1) using Seurat (V5.1.0) and merged. Cells with fewer than 200 detected genes and more than 5% mitochondrial reads were excluded from downstream analysis. The expression data was normalized using SCTransform and Principal Component Analysis (PCA) was performed. The reduced data was integrated by sample using Harmony (theta = 3, V3.8), clustered using FindNeighbors (dims=1:15) and FindClusters (resolution = 0.5). To visualise the clusters Uniform Manifold Approximation and Projection (UMAP) was performed and a UMAP generated using RunUMAP (spread = 2, min.dist = 0.4). A total of 19 clusters were identified, with the top markers for each cluster identified using FindAllMarkers (only.pos = TRUE, min.pct = 0.25, logfc.threshold = 0.25). Cluster annotation was performed manually based on differentially expressed markers (Table S1+ S2).

#### Analysis of scRNA-seq data

Significantly differentially expressed genes between treatments (adjusted P < 0.05 and log_2_ fold change > 0.5 or <(-0.5) were kept for further analysis. Pathway analysis was performed by the enrichGO function using the org.Mm.eg.db (version 3.20.0) database as reference.

### Statistical analysis

All statistical analyses were performed using GraphPad Prism software. Data are shown as mean and standard deviation. Data were tested for normality using the Shapiro-Wilk normality test. For comparisons with two variables, statistical differences were calculated using a two-way ANOVA with multiple comparisons. For comparisons with one variable, where data were normally distributed, a Student’s t-test was used for comparisons between two groups and a one-way ANOVA with Tukey’s multiple comparisons correction was used for comparisons between multiple groups. Where data were not normally distributed, a Mann Whitney U test (two groups) or a Kruskall-Wallis test with Dunn’s multiple comparisons correction (multiple groups) was used. Significance was determined based on *p*-value where \**p*< 0.05, \*\**p*<0.01, \*\*\**p*< 0.001, \*\*\*\**p*< .0001.

### Illustrations

All illustrations were created using BioRender.com.

## Abbreviations

cDC: conventional dendritic cell
cLP: colonic lamina propria
cMLN: colon draining mesenteric lymph node
DSS: dextran sulphate sodium
FACS: fluorescence activated cell sorting
IBD: inflammatory bowel disease
Hh: Helicobacter hepaticus
IL-10: interleukin-10
ILC: innate lymphoid cell
iNOS: inducible nitric oxide synthase
LPL: lamina propria leukocytes
LPS: lipopolysaccharide
mAb: monoclonal antibody
MHC: major histocompatibility complex
PAMP: pathogen associated molecular pattern
PRR: pattern recognition receptor
RT: room temperature
TLR: toll-like receptor
Treg: regulatory T cell
tSNE: t-distributed stochastic neighbour embedding
SCID: severe combined immunodeficiency
scRNA-seq: single cell RNA-sequencing
SFB: segmented filamentous bacteria
SPF: specific pathogen free
RAG: recombination activating gene
UMAP: uniform manifold approximation and projection

## Acknowledgements

We thank staff at the Central Research Facility (University of Glasgow) for animal husbandry; staff at the Cellular Analysis Facility (University of Glasgow) for support with flow cytometry, histology, and microscopy – in particular Diane Vaughan, Fiona McMonagle and Tyler Shaw; staff at the Molecular Analysis Facility (University of Glasgow) for their support with single cell RNA-sequencing; and Professor Fiona Powrie and Claire Pearson for providing the anti-mouse CD4 clone GK1.5 monoclonal antibody used in Figure 6. This work was funded by a Wellcome Trust Investigator Award (102972/Z/13/Z) to K.J.M, a MRC Project Grant (MR/X002004/1) to K.J.M. and the University of Glasgow through PhD studentship funding granted to K.J.M and A.L.L.H. A.A.M.A. is supported by the University of Glasgow Integrated Infection Biology PhD Programme from the Wellcome Trust (218518/Z/19/Z).

## Author contributions

A.L.L.H. designed and performed experiments, analysed data, produced figures, and wrote the manuscript. A.A.M.A. performed experiments, analysed data, produced figures, and wrote the manuscript. A.F. performed experiments and analysed data, X.L. performed experiments. D.M. and J.C. performed processing and bioinformatics analysis of sequencing data. T.R.T. performed experiments. H.C.W. designed and performed preliminary experiments. K.J.M. conceptualised the study, obtained funding, supervised the project, designed experiments, and wrote the manuscript.

