## Supplementary Figures for "The murine intestinal pathobiont *Helicobacter hepaticus* attenuates DSS colitis in a CD4+ T cell-dependent manner"

### Supplementary figure legends

#### Supplementary Figure 1: Flow cytometric analysis of CD45$^+$ cLP immune populations

(A) Gating strategy for identification of cLP myeloid populations by flow cytometry.

(B) Gating strategy for identification of cLP lymphoid populations by flow cytometry.

(C) tSNE clustering and feature plots showing expression of flow cytometry markers across myeloid populations.

(D) tSNE clustering and feature plots showing expression of flow cytometry markers across lymphoid populations.

#### Supplementary Figure 2: Changes to colonic immune populations and cytokine transcription following *Hh* colonization and DSS colitis

C57BL/6 littermates were infected with $1x10^8$ CFU *H. hepaticus* or given PBS by oral gavage and administered 2% DSS in drinking water from days 21-24 or given normal water as controls. Mice were culled on day 29.

(A) Flow cytometric analysis of $CD45^+$ colonic LPL populations.

(B) A section of distal colon was isolated and expression of indicated genes measured using qPCR. Differences in gene expression were determined using the $2^{-\Delta\Delta C(t)}$ method, with gene expression normalised to the housekeeping gene *Rps29* and data shown as fold change relative to the PBS control group.

Data are presented with mean ± SD and are pooled from 3 independent experiments with n=4-6 per experiment. Each data point represents an individual mouse. Statistical significance was determined by 2-way ANOVA (significance *p< 0.05, **p< 0.001, ***p< 0.001, ****p< .0001).

#### Supplementary Figure 3: *In vivo* treatment with αTLR2 mAb inhibits responses to Pam3Csk4 and does not affect *Hh* colonization levels

(A) C57BL/6 littermates received either 100µg αTLR2 mAb or mouse IgG1, κ isotype control mAb by i.p. injection. 4 days later, mice were challenged with 100µg Pam3csk4 or received PBS by i.p. injection. Mice were culled after 4 hours and concentrations of IL-6 and MCP-1 measured in the serum by CBA.

(B) C57BL/6 littermates were infected with $1x10^8$ CFU *Hh* or given PBS by oral gavage. All groups were administered 2% DSS in drinking water from days 21-24 before being culled on day 29. Mice received either 100µg αTLR2 mAb or mouse IgG1, κ isotype control mAb by i.p. injection once weekly on days 0, 7, 14 and 21. *Hh* DNA was quantified in caecal contents on day 29 by qPCR of the *Hh*-specific *CdtB* gene.

Data are presented with mean ± SD and represent one experiment (A) or are pooled from 2 independent experiments with n=5 per experiment (B). Each data point represents an individual mouse. Statistical significance was determined by 2-way ANOVA (A) or Mann-Whitney test (B) (significance *p< 0.05, **p< 0.001, ***p< 0.001, ****p< .0001).

#### Supplementary Figure 4: Expression of Foxp3 and RORγt are not altered on *Hh*-specific T cells during DSS colitis

C57BL/6 littermates were infected with $1x10^8$ CFU *Hh* and administered 2% DSS in drinking water from days 21-24 or given normal water as controls. Mice were culled on day 29. *Hh* specific $CD4^+$ T cells were identified using a fluorescently conjugated HH1713 tetramer. Representative dot plots of cLP and cMLN HH1713 $tetramer^+$ ($Tet^+$) $CD4^+$ T cells, analysed for expression of Foxp3 and RORγt (upper) and quantification (lower). Data are presented with mean ± SD and are pooled from 3 independent experiments with n=3-6 per experiment. Each data point represents an individual mouse. Statistical significance was determined by 2-way ANOVA (significance *p< 0.05, **p< 0.001, ***p< 0.001, ****p< .0001).

#### Supplementary Figure 5: Changes to cLP $CD4^{+}$ T cell phenotypes following streptomycin-induced clearance of *Hh* colonization

C57BL/6 littermates were infected with $1x10^{8}$ CFU *Hh* or given PBS by oral gavage, before certain groups received 20mg streptomycin by i.p. injection on days 21 and 22. Streptomycin treated mice were then co-housed with an experimentally naïve littermate for 2 weeks to promote microbiota normalization following antibiotic treatment. Naïve littermates were then removed and experimental mice administered 2% DSS in drinking water from days 35-38 or given normal water as controls. Mice were culled on day 43.

(A-F) T cell numbers were quantified in the cLP and cMLN on day 43 using flow cytometry.

(A) $CD4^{+}$ T cells

(B) $Foxp3^{-}$ $ROR\gamma t^{+}$ of $CD4^{+}$ T cells

(C) $Foxp3^{+}$ $ROR\gamma t^{-}$ of $CD4^{+}$ T cells

(D) $Foxp3^{+}$ $ROR\gamma t^{+}$ of $CD4^{+}$ T cells

(E) $ROR\gamma t^{-}$ $Tbet^{+}$ of $Foxp3^{-}$

(F) $ROR\gamma t^{+}$ $Tbet^{+}$ of $Foxp3^{-}$

Data are presented with mean ± SD and are pooled from 3 independent experiments with n=3-6 per experiment. Each data point represents an individual mouse. Statistical significance was determined by 2-way ANOVA (significance *$p< 0.05$, **$p< 0.001$, ***$p< 0.001$, ****$p< .0001$).

#### Supplementary Figure 6: Cluster annotation of cLP CD3$^+$ T cells analyzed by scRNA-sequencing

C57BL/6 littermates were infected with 1x10$^8$ CFU *H. hepaticus* or given PBS by oral gavage and culled at day 21, total CD3$^+$ T cells were sorted from cLP and underwent scRNA-sequencing.

A) UMAP visualization of total T cell subsets in lamina propria in merged naïve and *Hh* samples with frequency distribution

B) Feature plots of relative expression data overlayed on UMAP visualization of T cell subsets for selected markers.

C) Heatmap showing expression of top 6 differentially expressed genes for each cluster when comparing clusters 1 to 18 in merged naïve and *Hh* samples, ranked by fold change

D) Volcano plot of differentially expressed genes in cluster 6 (tTreg), cluster 9 (pTreg_2), cluster 7 (Th1_2) and cluster 12 (Proliferating CD4 T cells.

### Supplementary materials

#### Supplementary Figure 1

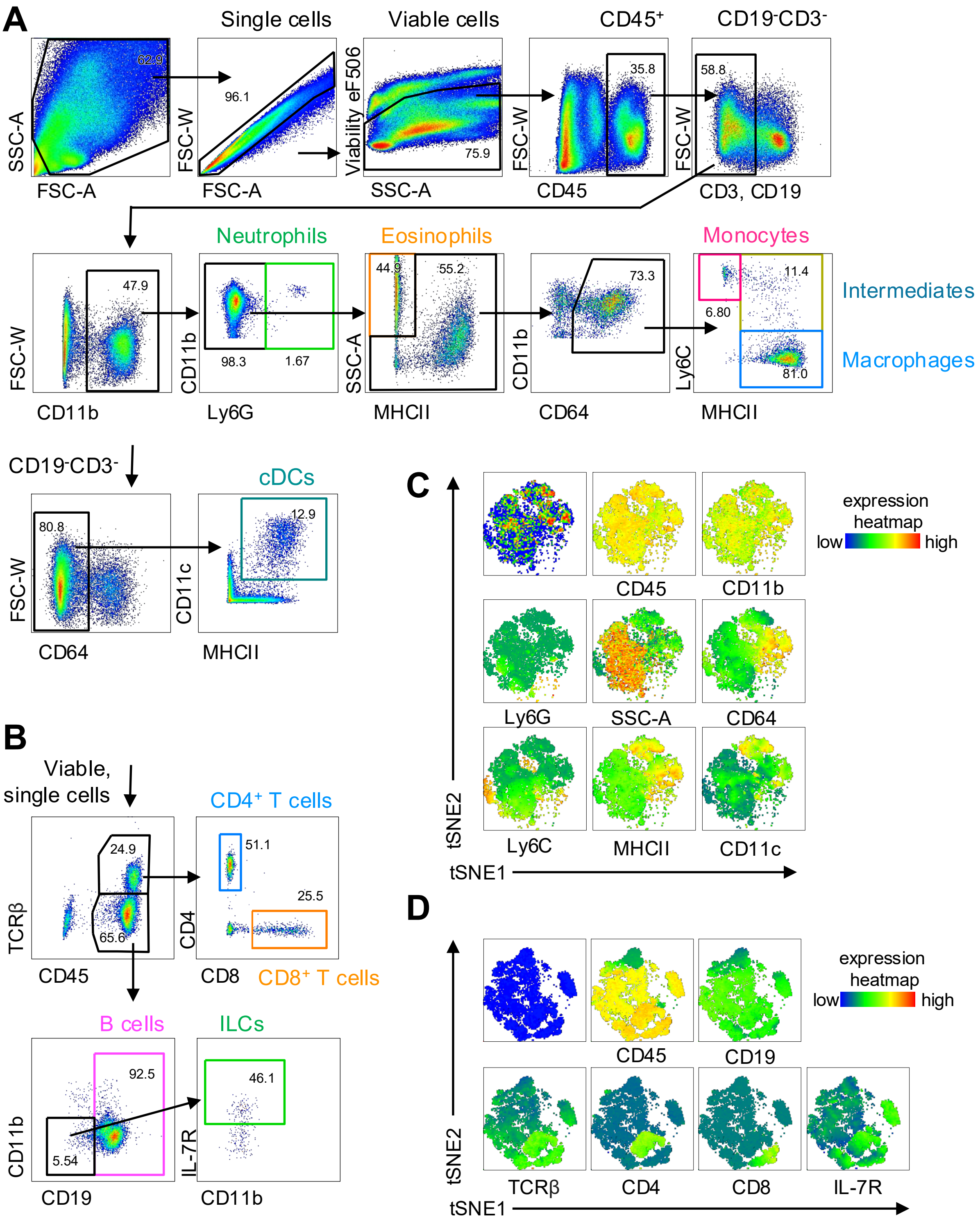

### Supplementary Figure 2

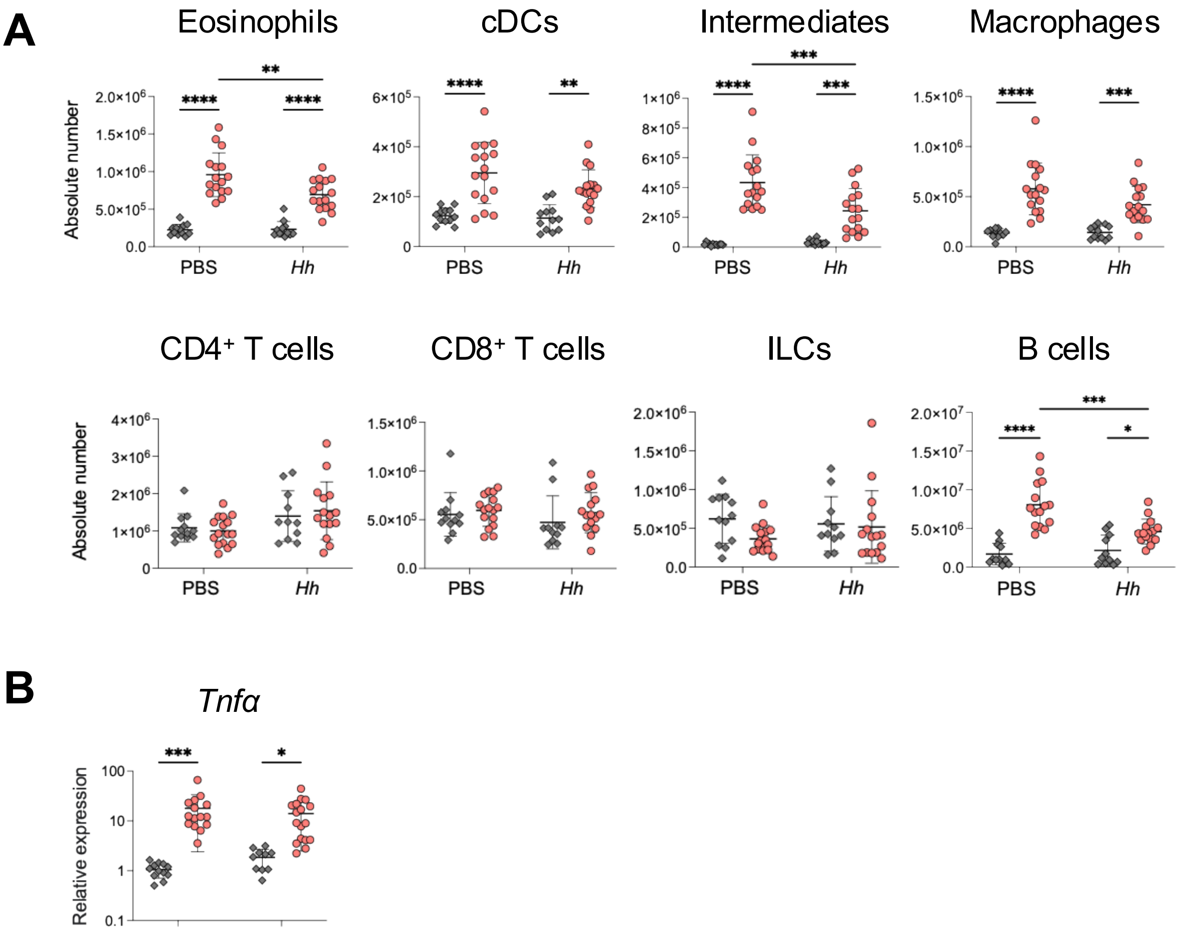

#### Supplementary Figure 3

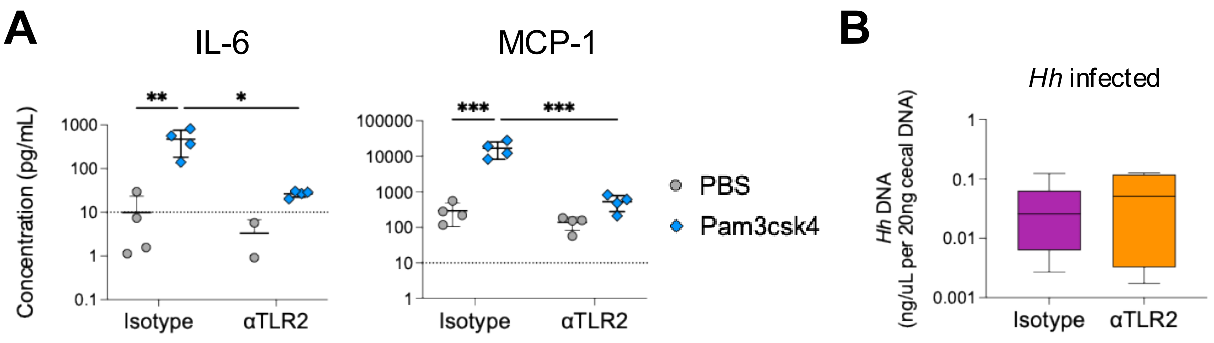

### Supplementary Figure 4

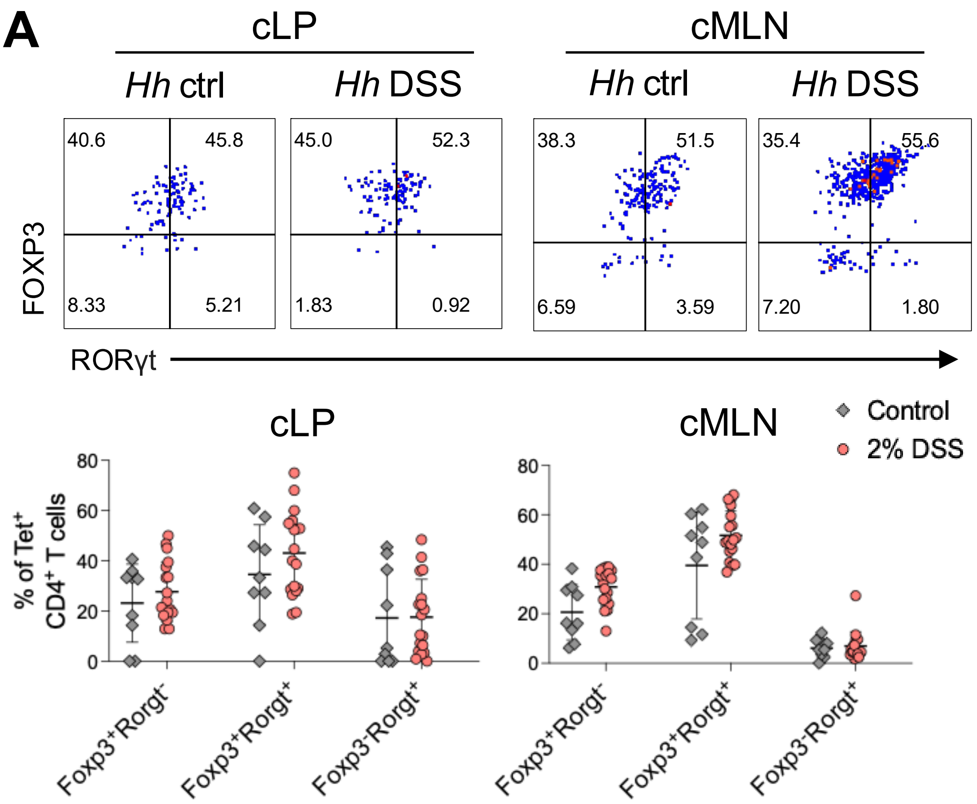

#### Supplementary Figure 5

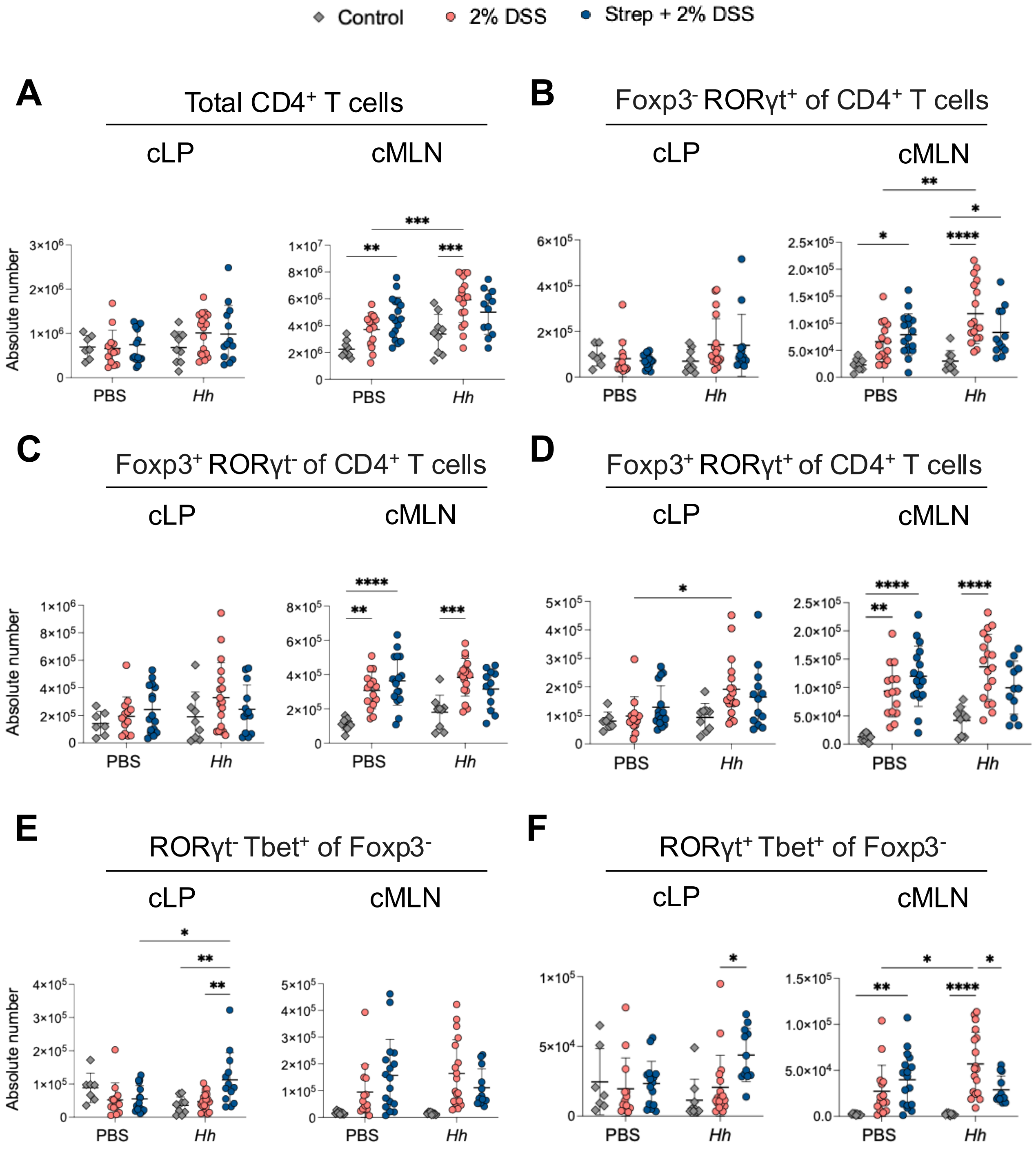

### Supplementary Figure 6

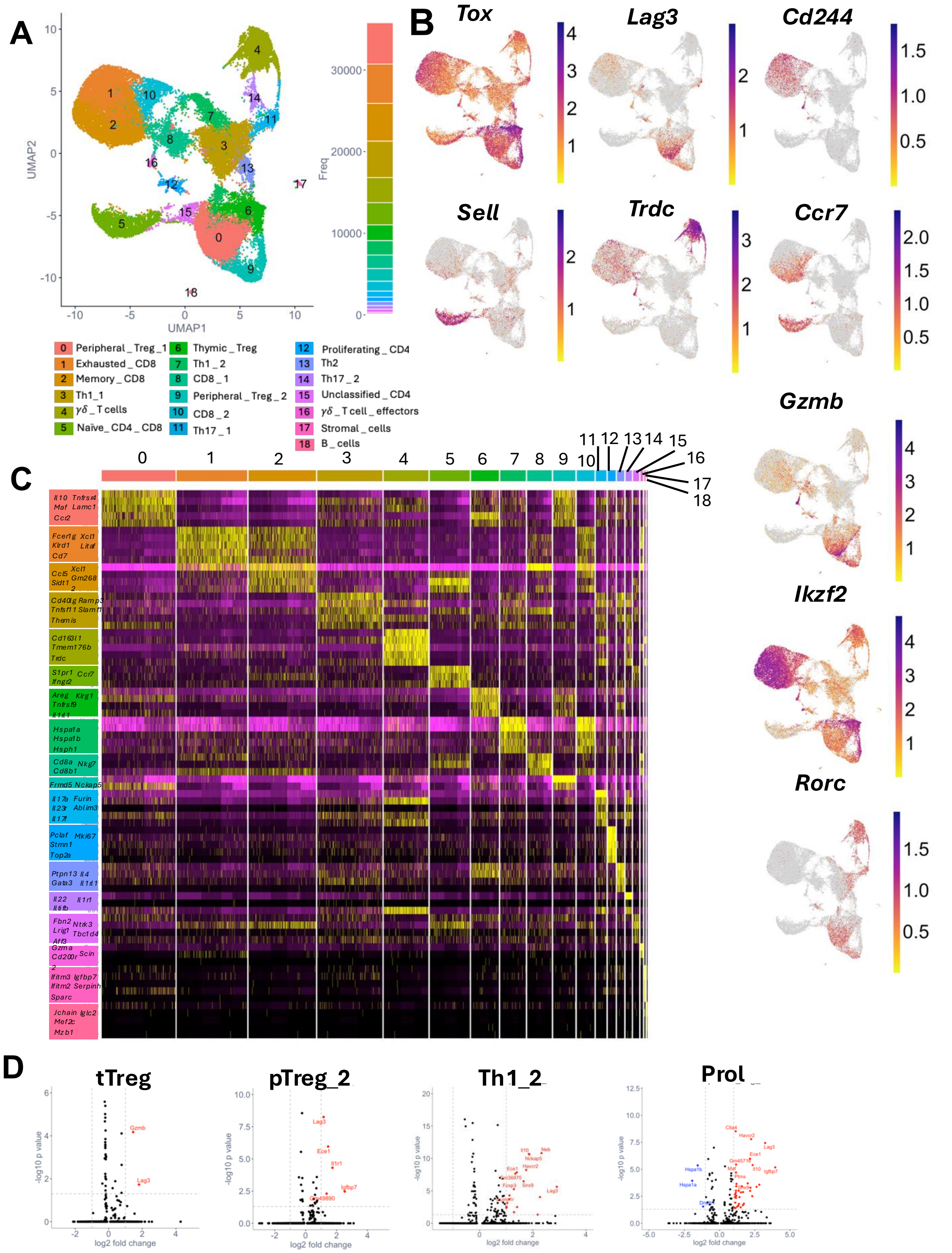

Supplementary Table 1: Top 10 defining gene markers per cluster and additional markers used for cluster annotation in scRNA-seq analysis

| Cluster | 0 | 1 | 2 | 3 | 4 | 5 | 6 | 7 | 8 | 9 | 10 | 11 | 12 | 13 | 14 | 15 | 16 | 17 | 18 |
|---|---|---|---|---|---|---|---|---|---|---|---|---|---|---|---|---|---|---|---|
| Top 10 markers | Il10 | Fcer1g | Sidt1 | Cd40lg | Cd163l1 | Ccr7 | Areg | Hspa1a | Cd8a | Frmd5 | Hspa1a | Il17a | Pclaf | Ptpn13 | Il22 | Lrig1 | Gzma | Col3a1 | Jchain |
| | Foxp3 | Cd7 | Klhdc2 | Tnfsf11 | Trdc | Lef1 | Tnfrsf4 | Hspa1b | Cd8b1 | Il10 | Hspa1b | Il17f | Birc5 | Il4 | Il23r | Aff3 | Cd200r2 | Postn | Mzb1 |
| | Nckap5 | Xcl1 | Ly6c2 | Themis | Zbtb16 | S1pr1 | Tnfrsf9 | Hsph1 | Ccl5 | Nckap5 | Hsph1 | Furin | Ccna2 | Pard3 | Il1r1 | Ntrk3 | Glp1r | Col6a1 | Gm30211 |
| | Neb | Cd160 | Pde2a | Ramp3 | Trdv4 | Ifngr2 | Il1rl1 | Dnajb1 | Ccl4 | Foxp3 | Dnajb1 | Ablim3 | Ube2c | Apbb2 | Tmem176a | Tbc1d4 | Dapk2 | Adamdec1 | Iglc2 |
| | Snx9 | Lrmda | Syde2 | Abcb1b | Ppp1r14c | Cmah | Klrg1 | Hsp90aa1 | Ifng | Neb | Rbpms | Tnfsf8 | Hmmr | 1700061F12Rik | Il17a | Izumo1r | Gzmk | Pcsk5 | Iglv1 |
| | Havcr2 | Tyrobp | Tsc22d1 | Btla | Rbpms2 | Igfbp4 | Cd83 | Bag3 | Gm44174 | Klrg1 | Id3 | Hlf | Esco2 | Gm42620 | Pxdc1 | Sh2d1a | Gzmb | Sfrp1 | Iglc1 |
| | Myo3b | Klra7 | Sox5 | St8sia1 | Klrb1b | Sell | Hopx | Dnaja4 | Plac8 | Ky | Rhob | Ccr6 | Pbk | Ccl1 | Ablim3 | Myb | Ppp2r2c | Pde3a | Ly6d |
| | Lag3 | Car2 | 1700025G04Rik | Abhd17c | Abi3bp | Cd8b1 | Nav2 | Hspd1 | Fasl | Msrb3 | Ly6c2 | Zeb2 | Ska1 | Il17rb | Pde5a | Il6ra | Tspan9 | Mfap4 | Tnfrsf17 |
| | Gzmb | Klra6 | Sell | Dipk2a | Blk | Rflnb | Cd81 | Ifng | Slamf7 | Myo3b | Klre1 | Kcnmb4 | Hist1h2ab | Calca | Ccr6 | St8sia1 | 2900026A02Rik | Bmp5 | Igha |
| | Sdcbp2 | Dapk2 | Xdh | Tnfsf8 | Zfp462 | Dapl1 | Ccr8 | Atf3 | Klrc1 | Il1rl1 | Bag3 | Ece1 | Hist1h3c | Itga2 | Kcnmb4 | Ap1s3 | Ccl3 | Abca8a | Igkc |
| Additional markers | Rorc | Cd8a | Cd8a | Tbx21 | Tcrd | | Foxp3 | Tbx21 | | Rorc | Cd8a | Rorc | Mki67 | Gata3 | Rorc | | Tcrd | | |
| | | Cd8b1 | Cd8b1 | Ifng | | | Il10 | Tnf | | | Cd8b1 | | | | | | Tcrg | | |
| | | Lag3 | Ccr7 | Tnf | | | Helios | | | | | | | | | | | | |
| | | Cd244 | Ccl5 | | | | | | | | | | | | | | | | |

#### Supplementary Table 2: Full cluster names for $CD3^{+}$ T cells identified by scRNA-seq

| Cluster | Name |
|---|---|
| 0 | Peripheral regulatory T cells_1 |
| 1 | Exhausted CD8 |
| 2 | Recently activated CD8 |
| 3 | Th1_1 |
| 4 | $\gamma\delta$ T cells |
| 5 | Naïve_CD4_CD8 |
| 6 | Thymically derived regulatory T cells |
| 7 | Th1_2 |
| 8 | CD8_effector_1 |
| 9 | Peripheral regulatory T cells_2 |
| 10 | CD8_effector_2 |
| 11 | Th17_1 |
| 12 | Proliferating CD4 |
| 13 | Th2 |
| 14 | Th17_2 |
| 15 | Unclassified CD4 |
| 16 | $\gamma\delta$ T cell effectors |
| 17 | Stromal contamination |
| 18 | B cell contamination |
